# The activation of *Chlamydomonas reinhardtii* alpha amylase 2 by glutamine requires its N-terminal ACT domain

**DOI:** 10.1101/2024.03.07.583849

**Authors:** Lisa Scholtysek, Ansgar Poetsch, Eckhard Hofmann, Anja Hemschemeier

## Abstract

The coordination of assimilation pathways for all the elements that make up cellular components is a vital task for every organism. Integrating the assimilation and use of carbon (C) and nitrogen (N) is of particular importance because of the high cellular abundance of these elements. Starch is one of the most important storage polymers of photosynthetic organisms, and a complex regulatory network ensures that biosynthesis and degradation of starch are coordinated with photosynthetic activity and growth. Here, we analyzed three starch metabolism enzymes of *Chlamydomonas reinhardtii* that we captured by a cyclic guanosine monophosphate-(cGMP-) affinity chromatography approach, namely soluble starch synthase STA3, starch branching enzyme SBE1 and α-amylase AMA2. While none of the recombinant enzymes was directly affected by the presence of cGMP or other nucleotides, suggesting an indirect binding to cGMP, AMA2 activity was stimulated in the presence of L-glutamine (Gln). This activating effect required the enzyme’s N-terminal aspartate kinase–chorismate mutase– tyrA (ACT) domain. Gln is the first N assimilation product and not only a central compound for the biosynthesis of N-containing molecules, but also a recognized signaling molecule for the N status. Our observation suggests that AMA2 might be a means to coordinate N- and C metabolism at the enzymatic level, increasing the liberation of C-skeletons from starch when high Gln levels signal an abundance of assimilated N.

## Introduction

Organisms need to acclimate to external or internal fluctuations in nutrient availability to adjust the biosynthesis of cell components that are mostly composed of more than one element. Carbon (C) and nitrogen (N) are the most abundant elements in cells, and their assimilation and distribution are tightly regulated to maximize growth. Plants and algae acquire N mostly through the assimilation of nitrate, nitrite or ammonium (Liu et al., 2022; Calatrava et al., 2023). Ammonium, the product of most N assimilation pathways, is ultimately transferred to L-glutamate (Glu) by glutamine synthetase (GS), forming L-glutamine (Gln). The Glu acceptor molecule is recycled by the transfer of the amino group to α-ketoglutarate (αKG; or 2-oxoglutarate) by glutamate synthase (also termed glutamine oxoglutarate aminotransferase, GOGAT), and the resulting net Glu is employed as an amino group donor for subsequent biosynthetic pathways.

In photosynthetic organisms, C is mostly acquired through carbon dioxide (CO_2)_ assimilation by the Calvin-Benson-Bassham cycle (CBB cycle). Triose phosphates and hexoses thus produced can be employed for anabolic processes directly or stored as starch, an insoluble glucose polymer, and one of the most important storage polymers in the Archaeplastida. Transitory starch is formed during the daily illumination period and degraded regularly, mostly in darkness, to provide C skeletons and energy for growth and maintenance, whereas storage starch serves as a mid-to long-term resource for periods of starvation, seedling development or perennation. The importance of coordinating starch metabolism with growth and survival is reflected by multiple regulatory interventions in starch biosynthesis and degradation, many of which have been discovered rather recently (Geigenberger, 2011; Santelia et al., 2015; Abt and Zeeman, 2020).

Starch consists of two polyglucan subfractions, which are both α-1,4-linked glucose polymers branched through α-1,6 glycosidic bonds (Bertoft, 2017). The major subfraction is amylopectin, which is regularly branched and forms the semicrystalline backbone of starch. Amylose, the minor fraction, usually consists of shorter glucose chains and only few branches. The enzymatic steps of starch biosynthesis and degradation are quite well understood (Zeeman et al., 2010; Deschamps et al., 2023). The glucose substrate for starch biosynthesis derives from fructose-6-phosphate, an intermediate of the CBB cycle, by the actions of phosphoglucose isomerase and phosphoglucomutase. The first committed substrate for starch biosynthesis is adenosine diphosphate-(ADP-) glucose, produced by ADP-glucose pyrophosphorylase (AGPase) from adenosine triphosphate (ATP) and glucose-1-phosphate. ADP-glucose is used by starch synthases to elongate glucan chains through α-1,4-glycosidic bonds, releasing ADP upon catalysis. Starch synthases belong to at least six different classes, soluble starch synthases SSI to SSV (although SSV is catalytically inactive) and granule-bound starch synthase (GBSS), and enzymes belonging to different classes have distinct roles in starch biosynthesis (Schwarte et al., 2013; Abt and Zeeman, 2020; Irshad et al., 2021; Deschamps et al., 2023).

Branches are introduced into the polyglucan by starch branching enzymes (SBEs) that cleave an α-1,4 glycosidic bond and form a new α-1,6 bond between the cleaved glucan and a glucose molecule of a linear chain. SBEs are grouped into class I and class II, and analyses of mutant plants and algae showed that the loss of class II branching enzymes usually results in notable phenotypes, whereas the absence of class I branching enzymes appears to have little or no consequences (Tetlow and Emes, 2014; Courseaux et al., 2023, and references therein). Two types of debranching enzymes – isoamylases and limit-dextrinases (also termed pullulanases) hydrolyze α-1,6 bonds (Zeeman et al., 2010; Deschamps et al., 2023). In the direction of starch granule formation, debranching enzymes ‘trim’ the branches of amylopectin so that the highly ordered structure of starch can be formed. Disproportionating enzymes or α-1,4 glucanotransferases transfer maltooligosaccharides from a donor to an acceptor glucan. In the unicellular green alga *Chlamydomonas reinhardtii* (*Chlamydomonas* henceforth), their role appears to be a ‘rescue’ of glucan chains released upon trimming (Wattebled et al., 2003, and references therein), but in the Brassicacean model plant *Arabidopsis thaliana* (*Arabidopsis* henceforth), they seem to be involved in starch degradation (Critchley et al., 2001; Li et al., 2017). Likewise, starch phosphorylases that reversibly catalyze the phosphorolytic cleavage of α-1,4-glucosidic bonds in linear chains, releasing glucose-1-phosphate, appear to have roles both in starch biosynthesis and degradation (Pfister and Zeeman, 2016).

For degradation, the insoluble starch granule must be made accessible to hydrolytic enzymes (Zeeman et al., 2010; Stitt and Zeeman, 2012). This is achieved by the phosphorylation of glycosyl residues by α-glucan, water dikinases and phosphoglucan, water dikinases. Amylases, debranching enzymes and starch phosphatases act in concert to hydrolyze the polyglucan chains: Beta-amylases are maltose-liberating exoamylases that require the phosphate groups at glucose moieties to be removed by starch phosphatases. Debranching enzymes remove branches that inhibit β-amylase activity. Alpha-amylases hydrolyze endogenous α-1,4-glycosidic bonds, and their importance for starch degradation appears to vary depending on the species. In *Arabidopsis*, β-amylases are the major starch hydrolyzing enzymes (Yu et al., 2005), whereas α-amylase I-1 is required for normal starch degradation in rice (Asatsuma et al., 2005). In *Chlamydomonas*, predominating α-amylase activity in partially purified amylase preparations was suggested (Levi and Gibbs, 1984). However, a *Chlamydomonas* mutant with a disruption of the β-amylase gene *AMB1* showed clear defects in storage starch degradation (Tunçay et al., 2013).

Starch synthesis and degradation are regulated on multiple levels that sometimes converge on one enzyme. For most catalytic steps, several enzyme isoforms are encoded by plant and algal genomes that differ in catalytic characteristics and/or domain architectures (Deschamps et al., 2023). The starch synthases of different classes, although in most cases sharing the catalytic domain, have diverse N-termini (Schwarte et al., 2013; Qu et al., 2018; Irshad et al., 2021), and the N-terminal coiled-coil domain of *Arabidopsis* SS4, for example, is required for correct localization and starch granule initiation (Raynaud et al., 2016; Lu et al., 2018). Differential gene expression also plays a role (e.g., Smith et al., 2004; Qu et al., 2018), but many regulatory signals work directly on the enzymes. Depending on plant species and localization, the first committed enzyme, AGPase, can be allosterically activated by 3-phosphoglycerate (3PGA), the product of CO_2 f_ixation by ribulose-1,5-bisphosphate carboxylase/oxygenase (Rubisco), and inhibited by phosphate (Geigenberger, 2011; Figueroa et al., 2022). Starch phosphorylases of several species, including *Chlamydomonas*, are inhibited by ADP-glucose (Dauvillée et al., 2006; Hwang et al., 2010). A number of enzymes of the starch metabolism are regulated through reversible disulfide bond formation (Kötting et al., 2010; Santelia et al., 2015; Skryhan et al., 2018). For example, AGPase activity and sensitivity to 3PGA are increased by the reduction of specific cysteine thiol groups, whereas the oxidative formation of a disulfide bond has the opposite effect (Geigenberger, 2011; Figueroa et al., 2022). Additional examples are the *Arabidopsis* β-amylase BAM1 (Sparla et al., 2006; Valerio et al., 2011) and α-amylase AMY3 (Seung et al., 2013) or potato α-glucan water dikinase (Mikkelsen et al., 2005) that are active in their reduced, but hardly active in their oxidized form.

Starch metabolism enzymes are also modified by protein phosphorylation (Kötting et al., 2010), which affects protein activity, for example in case of wheat SBEIIa and SBEIIb, as well as protein complex formation (Tetlow et al., 2004; Mehrpouyan et al., 2021). Indeed, enzymes of the starch metabolism have been detected in (multi-) protein complexes, suggesting a concerted action of these enzymes or regulatory interactions (Kötting et al., 2010; Geigenberger, 2011). In some cases, protein complexes are formed between catalytically active enzymes and inactive isoforms, or with dedicated scaffolding or targeting proteins (Abt and Zeeman, 2020). One example is *Arabidopsis* LIKE SEX4 1 (LSF1), an inactive starch phosphatase that interacts with plastidial β-amylases and is required for normal starch degradation (Schreier et al., 2019).

Previously, we showed that the *Chlamydomonas* nitric oxide-(NO-) sensitive guanylate cyclase CYG12, which we had suspected to be involved in hypoxic acclimation, apparently plays pleiotropic roles (Düner et al., 2018): An algal strain in which the *CYG12* transcript was post-transcriptionally down-regulated by a micro-RNA approach showed impaired growth in hypoxia and darkness, and additionally photosynthetic defects in aerobiosis in the light. The strain also exhibited aberrant patterns of consuming acetate from the medium and degrading internal starch reserves. Here, we show the results of an affinity chromatography approach that aimed to capture proteins that bind to cyclic guanosine monophosphate (cGMP), the product of guanylate cyclase activity. With this technique we aimed to identify possible targets of CYG12 signaling. We were surprised that, besides identifying several proteins predicted to bind cyclic nucleotide monophosphates (cNMPs), we found 15 proteins that are involved in the metabolism of starch. We selected three enzymes for biochemical analyses, soluble starch synthase 3 (STA3 or SSS3A), starch branching enzyme 1 (SBE1) and α-amylase 2 (AMA2). We focused on AMA2, because this protein has an N-terminal ACT (aspartate kinase – chorismate mutase – TyrA) domain, known to bind small ligands and thereupon regulating enzymatic activity (Grant, 2006; Lang et al., 2014). Notably, while none of the three selected enzymes was directly affected by cGMP or additional nucleotides *in vitro*, suggesting that they might bind to cGMP indirectly, we observed that AMA2 activity was increased in the presence of L-glutamine (Gln). This effect required the presence of the ACT domain and could be diminished by selected amino acid exchanges. This result suggests that within the cell, AMA2 might be important for the coordination of N- and C metabolism, being stimulated to release C skeletons when N is sufficiently available as indicated by its first assimilation product, Gln.

## Materials and methods

### cGMP interaction chromatography and protein identification

#### Preparation of *Chlamydomonas* soluble protein extracts

Soluble proteins were obtained from *Chlamydomonas* wild-type strain CC-124 mt^−^ [137c] (Chlamydomonas Resource Center, University of Minnesota, MN, USA). Cells were grown in liquid Tris acetate phosphate (TAP) medium (Harris, 1989) at 20°C upon continuous bottom-up illumination of 100 µmol of photons × m^-2^ × s^-1^ (two Osram Lumilux Cool White light bulbs alternating with one Osram Fluora bulb) on a reciprocal shaker set to 100 rpm. At a density of 3 × 10^6^ cells × mL^-1^, the cells were harvested by centrifugation (10 min, 3,200 *g*, 4°C). One gram of fresh weight was resuspended in 2 mL of extraction buffer (50 mM Tris-HCl, pH 7.4, 0.25 M sucrose, 1 mM EDTA, 0.1 mM MgSO_4,_ 10 mM KCl, 5 mM ascorbic acid, 1 mM phenylmethylsulfonyl fluoride, 1× protease inhibitor cocktail (cOmplete ULTRA tablets, EDTA-free, Roche), 0.5 % (w/v) polyvinylpolypyrrolidone (PVPP)). Cell lysis was done by sonication on ice (Branson Sonifier 250; output 40 %, 6 × 30 s with 90 s breaks). The lysate was centrifuged (30 min, 16,000 *g*, 4°C) and the supernatant was filtrated through a 0.2 µm syringe filter (cat-no. 83.1826.001; Sarstedt, www.sarstedt.com). The protein concentration of the filtered lysate was determined using the Bio-Rad Protein Assay Dye Reagent Concentrate (Bio-Rad; www.bio-rad.com) and bovine serum albumin (BSA) as standard and was subsequently adjusted to a concentration of 3 mg × mL^-1^ with extraction buffer.

**cGMP interaction chromatography** was done based on a protocol from Donaldson and Meier (2013) and Donaldson et al. (2016), respectively. All steps were performed in the cold room at 8°C. Two types of cGMP-functionalized agarose beads from the Biolog Life Science Institute (www.biolog.de) ((2’-AHC-cGMP-agarose (2’-O-(6-Aminohexylcarbamoyl)guanosine-3’, 5’-cyclic monophosphate-agarose; cat-no. A 059) and 2-AH-cGMP-agarose (N²-(6-Aminohexyl)guanosine-3’, 5’-cyclic monophosphate-agarose; cat-no. A 056)) as well as a control bead material functionalized with ethanolamine (EtOH-NH-agarose; cat-no. E 010) were prepared by mixing 200 µL of resuspended bead material with 1 mL of assay buffer (extraction buffer without PVPP) and incubating the mixture for 2 h on a shaker at 300 rpm. Afterwards, the agarose beads were harvested by mild centrifugation (30 s, 100 *g*) and the supernatants carefully removed. Each bead type was mixed with 1 mL of *Chlamydomonas* protein extract and incubated overnight under permanent shaking at 300 rpm. Following this incubation, the agarose beads were harvested by gentle centrifugation (30 s, 50 *g*) and the supernatant was removed. The beads were then resuspended in 1 mL of assay buffer, incubated for 5 min mixing at 300 rpm, and harvested by centrifugation. After discarding the supernatant, this washing step was repeated five times. Afterwards, nucleotides dissolved in assay buffer were employed to elute proteins with different nucleotide-binding characteristics, namely, in this order, 100 mM ADP, 100 mM GMP, 10 mM cyclic adenosine monophosphate (cAMP), 25 mM cAMP, 10 mM cGMP, and 100 mM cGMP. In each case, the agarose beads were resuspended in 200 µL of these solutions and incubated for 5 min at 300 rpm. After centrifugation, the supernatants were collected for subsequent protein identification. Each elution step was followed by a washing step as described above.

#### Preparation of tryptic peptides

For protein identification by mass spectrometry, the six elution fractions of each bead type were prepared for in-gel tryptic digest. First, proteins were precipitated by mixing each sample with 800 µL of –20°C cold acetone. After incubation overnight at –20 °C, the samples were centrifuged (10 min, 14,000 *g*, 4°C), the supernatant was discarded, and the protein pellets were briefly dried under a sterile bench. Then, the protein pellets were resuspended in 15 µL ultrapure water and 4 µL sodium dodecyl sulfate polyacrylamide gel electrophoresis-(SDS-PAGE) sample buffer (0.05 M Tris-HCl pH 8, 5 % (v/v) glycerol; 1.5 % (w/v) SDS; 0.05 mg × mL^-1^ bromophenol blue; 2.5 % (v/v) β-mercaptoethanol), incubated at 95°C for 5 min, and loaded on 15 % SDS polyacrylamide gels, separated by empty lanes. The agarose beads remaining after the final washing step were directly dissolved in SDS-PAGE sample buffer, heated, and loaded onto the gels. After the samples had run into the separating gel, electrophoresis was stopped, the gels were stained with Coomassie, and the protein containing parts were excised. The gel pieces were destained by first incubating them in 200 µL of 50 mM ammonium bicarbonate (ABC) for 20 min and mixing at 300 rpm, followed by several incubation steps in 25 mM ABC, 50 % (v/v) acetonitrile (ACN), until the gel pieces were colorless. The gel pieces were dehydrated in 100 % (v/v) ACN and then rehydrated with 30 µL of 3.33 µg × mL^-1^ trypsin (cat-no. T6567 from Sigma-Aldrich/Merck, www.sigmaaldrich.com) in 25 mM ABC. Afterwards, 25 mM ABC was added to fully cover the gel pieces, and tryptic digest was performed at 37°C overnight upon mixing at 300 rpm. After transferring the supernatant to a fresh reaction tube, 70 µL of 50 % (v/v) ACN, 1 % (v/v) formic acid (FA) were added to the gel pieces, followed by an incubation at room temperature and mixing at 300 rpm for 20 min. After centrifugation at 14,000 *g* and 4 °C for 1 min, the supernatant was combined with the first supernatant. This latter extraction step was repeated once. After the addition of 30 µL of 305 mM ammonium carbonate, 0.7 mM EDTA, the samples were completely dried in vacuum concentrator at room temperature. The pellets were frozen at –20 °C until further use.

**For mass spectrometric analyses** according to Cormann et al. (2016), the peptide pellets were resuspended in 20 µL of 0.1 (v/v) % FA, injected into a UPLC Symmetry C_18 t_rapping column (5 μm, 180 μm × 20 mm) and afterwards transferred to a UPLC BEH C_18 c_olumn (1.7 μm, 75 μm × 150 mm) using a nanoACQUITY UPLC system (all from Waters, www.waters.com). The temperature of the column oven was set to 45°C. Elution of peptides was achieved with a multistep gradient of buffer A (0.1% (v/v) FA) to buffer B (0.1 % FA in ACN) at a flow rate of 0.4 μL × min^-1^ (0 – 5 min: 1 % buffer B; 5 – 10 min: 5 % buffer B; 10 – 175 min: 40 % buffer B; 175 – 200 min: 99 % buffer B; 200 – 210 min: 1 % buffer B). Coupling to the mass spectrometer (Orbitrap Elite; Thermo Fisher Scientific, www.thermofisher.com) was achieved via a PicoTip Emitter (SilicaTip, 30 μm; New Objective, www.newobjective.com) and the spray voltage was at 1.5–1.8 kV. The temperature of the desolvation capillary was set to 200°C. Linear ion trap and Orbitrap were run in parallel during a full scan between 300 and 2,000 m/z and a resolution of 240,000 on the orbitrap. MS/MS spectra of the 20 most intense precursors were detected in the ion trap. For collision-induced dissociation, relative collision energy was set to 35 %, and dynamic exclusion was enabled (repeat count of one; 1-min exclusion duration window). Ions with an unassigned charge or singly or doubly charged ions were rejected.

The cGMP affinity chromatography experiment was done in two independent experiments, and computational protein identification was done on both sample sets simultaneously. The MaxQuant software (version 1.6.7.0) was used for data analysis, with all false discovery rates (FDRs) set to 0.01. Minimum peptide length was set to 6; maximal missed cleavages were set to 2; methionine oxidation and N-terminal acetylation were included as possible modifications. Decoy mode (revert) was enabled, and common protein contaminants were included. Protein identification was done against *C. reinhardtii* v5.6 proteins (Creinhardtii_281_v5.6.protein) to which the chloroplast proteome (derived from sequence FJ423446.1 at the European Nucleotide Archive; www.ebi.ac.uk/ena/) and mitochondrial protein sequences (from GenBank NC_001638) had been added manually.

### Heterologous production and purification of *Chlamydomonas* starch metabolism enzymes and variants

#### Generation of expression plasmids

The *Chlamydomonas* coding sequences for AMA2 (α-amylase; *Cre08.g362450.t1.*2; note that all gene identifiers refer to the *Chlamydomonas* genome version v5.6 at Phytozome 13), SBE1 (starch branching enzyme; *Cre06.g289850.t1.*1) and STA3 (soluble starch synthase III; *Cre06.g282000.t2.1*) were cloned into the *Escherichia coli* expression vector pASK-IBA7 (IBA Lifesciences GmbH; www.iba-lifesciences.com/). Expression from this vector equips the target protein with an N-terminal *Strep*-tactin affinity tag (*Strep*-tag II), followed by a Factor Xa cleavage site (Supporting Fig. S1). Oligonucleotides were designed using the Primer D’Singer 1.1 software (IBA Lifesciences GmbH) to amplify the coding sequences including the putative chloroplast targeting peptides (Table 1). In the case of AMA2, protein variant ΔAMA2 was additionally generated. It consists of the predicted amylase domain only and lacks the N-terminal 269 amino acids encompassing the putative chloroplast transit peptide, the predicted ACT (aspartate kinase – chorismate mutase – TyrA) domain as well as the sequence following the ACT domain up to the predicted amylase domain (Supporting Fig. S1). All sequences were amplified from total cDNA obtained from *Chlamydomonas* strain CC-124 mt^−^ as described in Huwald et al. (2015). Polymerase chain reactions (PCRs) were done employing the KAPA HiFi HotStart ReadyMix PCR Kit (#KK2601; Roche Sequencing, https://sequencing.roche.com/en.html). Sequences and vector were cut by *Bsa*I followed by ligation and transfection into *E. coli* DH5α by heat shock transformation. The cells were selected on LB agar plates (LB Lennox; Carl Roth GmbH, www.carlroth.com) containing 100 µg × mL^-1^ ampicillin, and colonies were subsequently grown in liquid LB Broth (Lennox) medium with 100 µg × mL^-1^ ampicillin. Plasmid isolation was done employing the GeneJet^TM^ Miniprep kit (ThermoFisher Scientific) and sequencing was performed by the DNA sequencing service at the chair for biochemistry, Biochemistry I, receptor biochemistry, at Ruhr University Bochum, Germany.

**Table 1.**
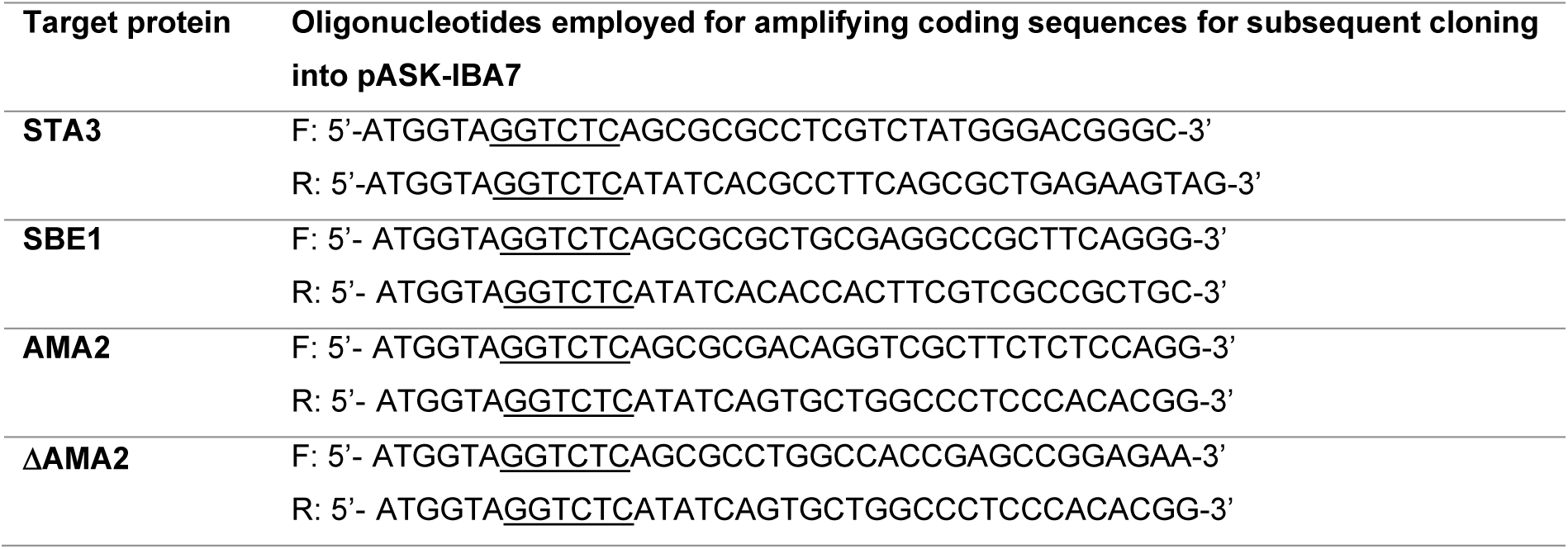
Oligonucleotides employed for amplifying the coding sequences of the selected enzymes. Oligonucleotides employed for amplifying the sequences coding for STA3 (Cre06.g282000.t2.1), SBE1 (Cre06.g289850.t1.1) and AMA2 (Cre08.g362450.t1.2) as well as AMA2 without the predicted ACT domain (ΔAMA2). *Bsa*I restriction sites (5′-GGTCTC(N_1_)/(N_5_)-3′), used for cloning into the expression vector pASK-IBA7, are underlined. F: forward, R: reverse.

#### Generation of AMA2 amino acid exchange variants

Single amino acids in the predicted ACT domain of AMA2 were exchanged by site-directed mutagenesis of the *AMA2* coding sequence within the pASK-IBA7 vector employing QuikChange PCR. Mismatched oligonucleotides where designed according to Zheng et al. (2004) and are listed in Table 2. Amplified PCR products were digested with *DpnI* and afterwards transfected into *E. coli* DH5α by heat shock transformation. Plasmid isolation and sequencing was done as described above.

**Table 2.**
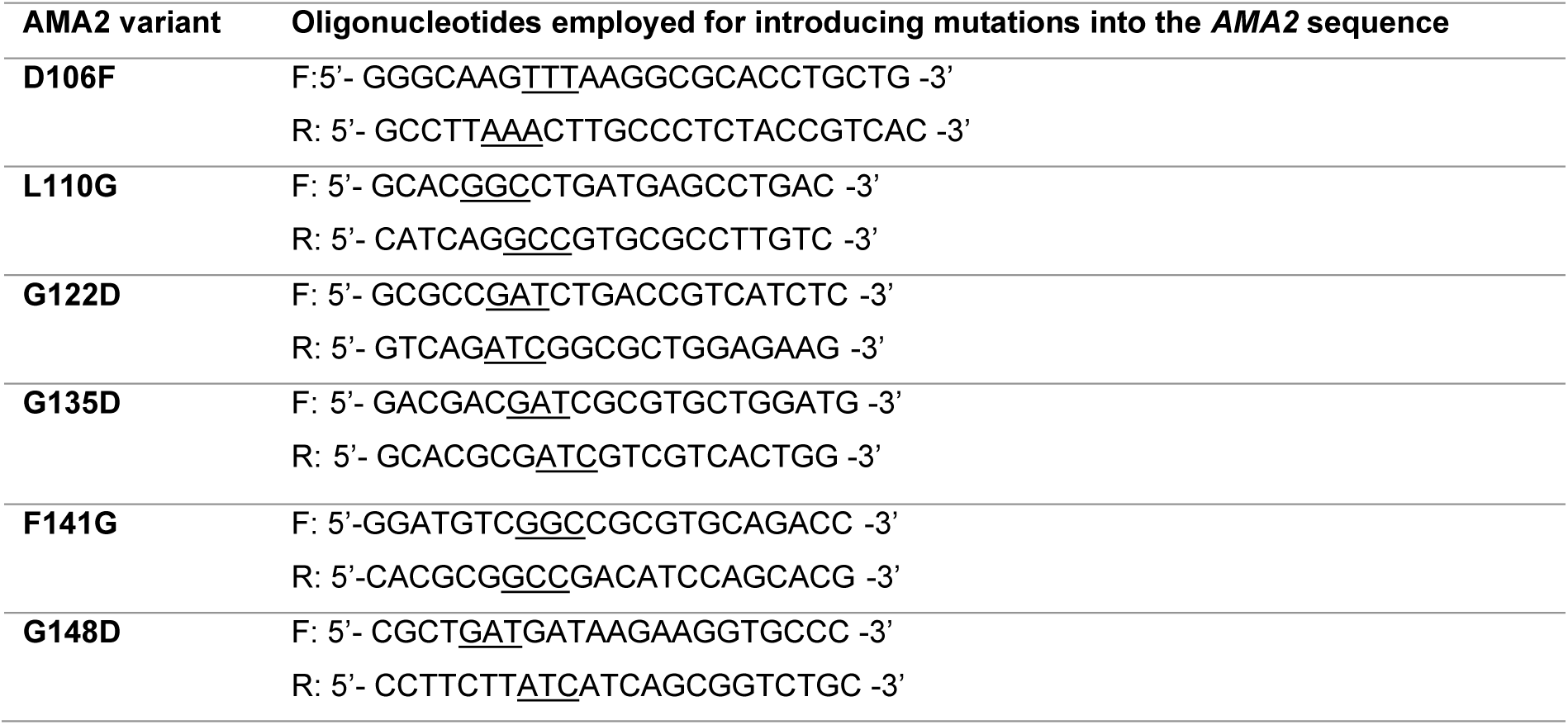
Oligonucleotides employed for generating AMA2 amino acid exchange variants. The codons that introduce mutations in the *AMA2* coding sequence are underlined. The numbers in column ‘AMA2 variant’ refer to the exchanged amino acid of the native protein including the N-terminal methionine residue.

**For heterologous protein production**, the expression vectors were introduced into *E. coli* Rosetta™ (DE3) by electroporation. Precultures were grown in 100 mL of LB medium containing 100 µg × mL^-1^ ampicillin and 25 µg × mL^-1^ chloramphenicol over night at 37°C, shaking at 120 rpm. Expression cultures were then inoculated to an OD_600 o_f 0.05 in 500 mL Terrific-Broth-(TB-) medium (12 g × L^-1^ tryptone, 24 g × L^-1^ yeast extract, 0.5 % (v/v) glycerol, 17 mM KH_2P_O_4,_ 72 mM K_2H_PO_4)_ supplemented with 2 mM MgCl_2,_ 100 µg × mL^-1^ ampicillin and 10 µg × mL^-1^ chloramphenicol in 2 L Erlenmeyer flasks. These cultures were incubated at 37°C, shaking at 120 rpm, until they had reached an OD_600 b_etween 0.5 and 0.6 (expression of *STA3* and *SBE1*) or between 0.9 and 1 (expression of *AMA2*). Expression of the constructs was then induced by the addition of 0.2 µg × mL^-1^ anhydrotetracycline from a stock solution of 2 mg × mL^-1^ in dimethyl sulfoxide, and the cultures were incubated overnight at 21°C, shaking at 120 rpm on a rotary shaker.

#### Protein purification

After about 17 h of expression, the *E. coli* cells were harvested by centrifugation for 20 min at 4,430 *g* at 4°C. The cell pellet was resuspended in Buffer W (100 mM Tris-HCl pH 8, 150 mM NaCl, 0.1 mM EDTA). Cells lysis was done by sonication on ice (output 50 %, 4 cycles á 45 s, 1 min breaks) with a Branson Sonifier 250 (Branson; www.emerson.com/en-us/automation/branson). The lysate was centrifuged at 180,000 *g* for 1 h at 4°C, and the supernatant was subsequently passed through a sterile filter (2 µm pore diameter). The cleared lysate was loaded onto a gravity-flow column with Strep-Tactin^®^ Superflow® high capacity resin (IBA Lifesciences GmbH) pre-equilibrated with Buffer W. Washing was done with 20 mL Buffer W, elution by applying 10 mL Buffer E (Buffer W with 2.5 mM desthiobiotin). Elution fractions were pooled, and proteins were concentrated using a 30 kDa Amicon Ultra-4 Centrifugal Filter Unit (Merck Milipore; www.merckmillipore.com). Protein concentration was determined employing the Bio-Rad Protein Assay Dye Reagent Concentrate (Bio-Rad) using BSA as standard. Size and purity of the proteins were routinely checked by separating protein solutions through SDS-PAGE.

#### Size exclusion chromatography (SEC)

SEC was performed to determine the oligomeric state of AMA2 and was done using ÄKTA protein purification systems and a Superdex 200 Increase 10/300 GL column (24 mL column volume). Buffer W was used for equilibration and elution. The flow rate was set to 0.75 mL × min^-1^ and proteins were loaded through a sample loop. Calibration of the column was performed using the Gel Filtration Markers Kit for Protein Molecular Weights 12,000-200,000 Da (#MWGF200; Sigma-Aldrich/Merck, www.sigmaaldrich.com).

### *In vitro* activity assays

#### Alpha-amylase assay

α-amylase activity was analyzed by recording the absorption of starch-iodine complexes adapted from the method of Xiao et al. (2006). Enzyme concentrations, buffers and starch types differed depending on the aim of the assay. In all cases, enzymes diluted in 200 µL of buffer were mixed with 200 µL of 0.7 mg × mL^-1^ starch dissolved in ultrapure water and then incubated shaking at 300 rpm for 30 min at 30°C except for experiments aimed to determine temperature optima. The reaction was stopped by adding 100 µL of 1 M HCl. The solutions were then mixed with 500 µL of iodine solution (0.52 mg × mL^-1^ I_2 a_nd 5.2 mg × mL^-1^ KI) and the absorption was determined at λ = 580 nm against 500 µL of buffer mixed with 500 µL of iodine solution. A control sample was prepared for each assay in the same way but without enzymes. Starch calibration curves were generated using 20 to 200 µg starch. Enzyme activity (U) was defined as the decrease of 1 mg of iodine-stainable starch per minute. Specific activity was calculated according to the formula U × µmol^-1^ = (A580_control –_ A580_sample)_ / ((A580 × mg starch^-1^) × 30 min × µmol enzyme)). Routinely, soluble potato starch was employed as substrate (cat.-no. S2630 from Sigma-Aldrich/Merck). However, we also tested enzyme activities on *Chlamydomonas* storage starch (see the procedure to isolate native starch below).

Three different buffers were employed depending on the purpose; the buffer type is always indicated in the text. Routine assays to check for the activities of freshly prepared enzyme batches were performed in 0.1 M potassium phosphate buffer, pH 7. In several cases, we used a buffer designed to approach standard cellular ion concentrations, which we termed ‘physiological buffer’ (50 mM potassium phosphate, pH 7.4, 50 mM KCl, 10 mM NaCl, 0.5 mM MgCl_2,_ 100 nM CaCl_2)_. For determining pH optima, 0.1 M Britton-Robinson Buffer, pH 5.0 to pH 11.0 (boric acid, phosphoric acid and acetic acid, titrated with sodium hydroxide to the desired pH), was employed (Britton and Robinson, 1931). Temperature optima were determined in Britton-Robinson Buffer, pH 7.5, at incubation temperatures between 10°C and 50°C.

#### Starch branching enzyme assay

SBE1 activity was determined as described for α-amylase above. In addition to starch as substrate, SBE1 activity was determined in the presence of amylopectin (amylopectin from maize; cat.-no. 10120 from Sigma-Aldrich/Merck). The assay was performed as described above, except that 200 µL of 0.9 mg × mL^-1^ amylopectin dissolved in ultrapure water were added. In this case, absorption was measured at λ = 550 nm. pH and temperature optima were determined as described for α-amylase, except that 0.1 M Britton-Robinson buffer, pH 7, was used for the latter.

#### Soluble starch synthase assay

Starch synthase activity was analyzed according to Kulichikhin et al. (2016) who established an assay in which ADP, released from ADP-glucose by starch synthase activity, is converted to NADPH by subsequent enzymatic reactions. First, STA3 dissolved in 200 µL water was added to a reaction mixture containing 50 mM HEPES-NaOH, pH 7.5, 1.4 mg amylopectin and 15 mM dithiothreitol (DTT) in a total volume of 423 µL. A reference tube was prepared in the same way. The mixtures were pre-incubated for 2 min at 30°C before adding 27 µL of 20 mM ADP-glucose. The reference tube was immediately incubated at 100°C for 1 min to stop the reaction, while the reaction tube was incubated for 20 min at 30°C before stopping the reaction at 100°C. For all subsequent steps, reference and experimental tubes were processed in parallel. After allowing the reaction mixtures to cool down to room temperature, the samples were centrifuged for 10 min at 10,000 *g*. 300 µL of the supernatants were transferred to fresh tubes, and 200 µL of solution 1 (50 mM HEPES-NaOH, pH 7.5, 200 mM KCl, 10 mM MgCl_2;_ 4 mM phosphoenolpyruvate (PEP)) as well as 1.2 U pyruvate kinase (P1506; Sigma-Aldrich/Merck) were added to convert ADP and PEP to ATP and pyruvate. The reaction mixtures were incubated for 20 min at 30°C, and the reactions were stopped by incubation at 100°C for 1 min followed by centrifugation. The mixtures were then transferred to new tubes containing 400 µL of solution 2 (50 mM HEPES-NaOH, pH 7.5, 10 mM glucose, 20 mM MgCl_2,_ 2 mM NADP^+^), and the reaction was started by adding 1.4 U hexokinase (H5000; Sigma-Aldrich/Merck) and 0.36 U glucose-6-phosphate dehydrogenase (G6PDH) (G7877; Sigma-Aldrich/Merck) to each tube. During this reaction, hexokinase employs ATP to form glucose-6-phosphate, which is oxidized by G6PDH, forming NADPH. The mixtures were incubated for 10 min at 30°C before the absorption of the reaction mixture was monitored at 340 nm against the reference mixture. The final absorbance value was recorded after 10 min. Enzyme activity (U) of soluble starch synthase was defined as the release of 1 µmol of ADP, which is equivalent to the production of 1 µmol of NADPH, per 1 min. The final amount of NADPH was calculated employing its specific absorption coefficient at λ = 340 nm (6.22 L × mmol^-1^ × cm^-1^) and related to the STA3 amount employed, taking into account all dilution steps. The pH optimum of STA3 was determined by replacing HEPES-by Britton-Robinson buffer, pH 5 to 12, in the first reaction mixture. Controls were performed when necessary to test for any effects on the additional enzymes present in this assay. In these cases, STA3 was excluded and 100 µM of ADP was added as substrate, whereas all other components and proceedings were the same as described above.

#### Effect of small molecules on enzyme activities

To test for any influence of small molecules on enzyme activities, assays were conducted as described above, employing our ‘physiological buffer’. The additional molecules were added in concentrations indicated in the results section. Proteinogenic amino acids, α-ketoglutarate, cGMP and cAMP, guanosine and adenosine monophosphate (GMP and AMP) were obtained from Sigma-Aldrich/Merck (www.sigmaaldrich.de). Cyclic diguanosine monophosphate (c-diGMP) and guanosine-3’,5’-bisdiphosphate (ppGpp) were obtained from Biolog Life Science Institute (C057; www.biolog.de/) and Jena Bioscience (NU-884; www.jenabioscience.com/), respectively.

#### Effect of redox agents

To test effects of reductants and oxidants, recombinant proteins were diluted to a concentration of 1 mg × mL^-1^ in 20 µL of ‘physiological buffer’. After the addition of 5 mM of DTT, or 0.1, 1 or 5 mM of diamide, the reaction mixtures were incubated for 20 min at 23°C. As a control, the same protein dilutions were incubated in parallel without the addition of redox reagents. Afterwards, *in vitro* activity assays were done as described above. Note that the first reaction mixture of the soluble starch synthae assay as described by Kulichikhin et al. (2016) routinely contains DTT, which was omitted for testing the effects of DTT and diamide on STA3.

### Purification of starch from *Chlamydomonas*

Native starch was purified from nitrogen-(N-) limited *Chlamydomonas* cultures by a protocol modified from Delrue et al. (1992). *Chlamydomonas* strain CC-124 was first grown in liquid TAP medium as described above. To induce N deficiency, the cells were inoculated at a density of 3 × 10^6^ cells × mL^-1^ in 250 mL of a modified TAP medium without any N source. After five days of growth, cells were harvest by centrifugation (6,000 *g* for 20 min at 4°C), the pellet was washed in 100 mL 20 mM HEPES-KOH, pH 7.5, and the cell suspension was transferred to 50 mL conical centrifugation tubes. After an additional centrifugation step, the cell pellet was resuspended in 2 mL of ice cold 20 mM HEPES-KOH, pH 7.5, and this suspension was diluted with 10 mL of lysis buffer (300 mM sorbitol, 50 mM HEPES-KOH, pH 7.5, 2 mM EDTA, 1 mM MgCl_2,_ 1 % (w/v) BSA). The algal cells were ruptured by sonication on ice (output 50 %, 6 cycles á 30 s, 1 min breaks), and then the lysate was distributed to 2 mL reaction tubes and centrifuged at 760 *g* for 10 min at 4°C. The supernatants were discarded, and the crude starch pellets were resuspended in 2 mL of lysis buffer. The suspensions were combined and carefully applied on a 24 mL Percoll gradient composed of three layers of 20 %, 45 % and 60 % Percoll in 0.9× lysis buffer. After centrifugation (15 min, 5,900 *g*, 4°C), the purified starch pellet was rinsed with 1 mL of lysis buffer and centrifuged at 2,000 *g* for 10 min at 4°C. The starch pellet was snap-frozen in liquid nitrogen and stored at –20°C. For *in vitro* enzyme assays, the starch pellet was thawed on ice and resuspended in 10 to 15 mL of distilled water. Starch concentration was analyzed by the starch-iodine method in different dilutions of the starch suspension, utilizing commercial soluble potato starch as a standard.

### *In silico* structure prediction and sequence analyses

The structure of an AMA2 dimer was predicted employing AlphaFold2-multimer as implemented in LocalColabfold (Jumper et al., 2021; Evans et al., 2022; Mirdita et al., 2022). Template information was applied, and no amber relaxation was used.

To generate alignments, we first extracted the AMA2 ACT domain sequence based on the AlphaFold2 models (also see results section). Residues 86 to 168 of the protein sequence Cre08.g362450.t1.2 were used as query in NCBI’s Basic Local Alignment Search Tool BlastP against the non-redundant protein sequences database. The hits thus retrieved were manually inspected to reduce the number of sequences. For example, many sequences from metagenomic annotations were omitted. Selected full-length sequences were analyzed by NCBI’s Conserved Domains or InterPro (Paysan-Lafosse et al., 2023) predictions to identify additional domains (Supporting Table S1). The aligned sequences as provided by the original NCBI BlastP analysis of the selected proteins were combined, and a few sequences were manually added. These were the two ACT domains of *Escherichia coli* GlnD (UniProt B6HZE1) (Zhang et al., 2010), the two ACT domains of *Arabidopsis thaliana* ACR11 (Sung et al., 2011) and the putative ACT domain of *Chlamydomonas* STA4 (Cre12.g552200.t1.2). In these three latter cases, the AlphaFold models deposited in the AlphaFold Protein Structure Database (https://alphafold.ebi.ac.uk/; GlnD: B6HZE1, ACR11: Q9FZ47, STA4: A8IYK1) were inspected to extract the residues encompassing the ACT domain. The final list of sequences was subsequently aligned using Clustal Omega (Sievers et al., 2011).

### Accession numbers

*Chlamydomonas* sequence data from this article can be found in the *Chlamydomonas reinhardtii* v5.6 genome annotation at Phytozome 13 (https://phytozome-next.jgi.doe.gov/info/Creinhardtii_v5_6) under the accession numbers indicated in the text and in Supporting Data Set 1.

### Supplemental Data files

Supporting Information (included in this file)

Supporting Data Set 1

## Results

### cGMP affinity chromatography captured starch metabolism enzymes

We employed cGMP affinity chromatography to identify downstream effectors of *Chlamydomonas* guanylate cyclases. This method has been applied for animal cyclic nucleotide monophosphate-(cNMP-) binding proteins (Kim and Park, 2003; Scholten et al., 2006) and adapted to *Arabidopsis* protein extracts (Donaldson and Meier, 2013). The procedure aims to enrich cGMP-binding proteins on agarose beads functionalized with cGMP, and sequential elution with ADP, GMP and cAMP reduces the number of unspecific proteins (Scholten et al., 2006). A control with agarose beads functionalized with ethanolamine helps to eliminate proteins that bind to the agarose matrix. The procedure employs non-denatured proteins, so that it captures proteins that bind directly, but also proteins that form complexes with proteins retained in the beads (Scholten et al., 2006; Scholten et al., 2007).

We conducted the cGMP affinity chromatography experiment twice in independent experiments, using independent *Chlamydomonas* protein extracts. Only proteins for which at least two unique peptides were detected in any given elution or agarose bead fraction were regarded as truly being present. In total, 1,246 *Chlamydomonas* proteins were detected (Supporting Data Set 1, Sheet 1). Of these, 319 proteins were present with maximally one unique peptide and were therefore not considered further. Of the remaining 927 proteins (Supporting Data Set 1, Sheet 2), we considered a protein as present in an elution fraction or retained in one of the bead types when they were represented by at least two unique peptides in both replicates. In case of the control bead fractions, however, we also had a look at proteins present only in one replicate to gain an idea about what types of proteins might be enriched by the bead material. 183 proteins were present with at least two unique peptides in both, and 223 in either of the control replicates (Supporting Data Set 1, Sheet 3). Many of these (135 proteins, 101 thereof in both replicates) were ribosomal proteins or translation factors. We noted that of the remaining proteins, many are predicted to bind nucleic acids, such as tRNA synthetases or DNA-dependent RNA polymerases (Supporting Data Set 1, Sheet 3).

We considered a protein as potentially (indirectly) cGMP-binding when it was detected with ≥ 2 unique peptides in the cGMP-functionalized agarose beads in replicate, but with maximally one unique peptide or an at least ten-fold lower intensity in the control bead replicates. Applying these constraints, 52 and 27 proteins were assigned as being retained in the 2-AH-cGMP- and 2’-AHC-cGMP agarose beads, respectively (Supporting Data Set 1, Sheet 4). Of these, 17 proteins were present in both bead types. We included in the list of possible cGMP binders also four proteins that were eluted by 100 mM cGMP from either bead type (Supporting Data Set 1, Sheet 4).

In the 2-AH-cGMP bead fractions, we detected four proteins that are predicted to contain canonical cNMP-binding domains, namely the Cyclic Nucleotide-Binding domain (CNB) of the prokaryotic catabolite gene activator (CAP) (or cAMP receptor protein; CRP) family (Rehmann et al., 2007) or the GAF-(c<u>G</u>MP-dependent phosphodiesterases (PDEs), <u>a</u>denylyl cyclases, and <u>F</u>hlA) domains (Aravind and Ponting, 1997; Heikaus et al., 2009). CNB domains are often found in cNMP-dependent protein kinases, whereas the GAF domain is present, for example, in cNMP-regulated PDEs. Of the proteins that we assigned as possible cGMP binders, three are annotated as protein kinases with several CNB domains (Cre02.g076900.t1.1 (FAP19); described in Wang et al. (2006), Cre12.g493250.t1.2 (FAP358) and Cre16.g663200.t1.1 (FAP295)) and one as a GAF domain-containing PDE (Cre13.g605100.t1.2 (PDE20)). None of these four proteins were also captured by the 2’-AHC-cGMP beads (Supporting Data Set 1, Sheet 4).

Of the remaining possible cGMP binders, many proteins are known or predicted to bind nucleotides such as ADP or GTP, nucleotide derivatives such as *S*-adenosylmethionine (SAM) and NAD(P)H or nucleic acids (Supporting Data Set 1, Sheet 4). We were surprised that we found 15 enzymes known or predicted to be involved in starch metabolism in our compilation, none of which has a canonical cNMP-binding domain, and most of which are not known to bind any nucleotides. All of these proteins were retained in the 2-AH-cGMP beads, and 10 of them in both beads, whereas none were only found in the 2’-AHC-cGMP beads.

We selected three enzymes from our list of possible cGMP binders to test for any effects this second messenger might exert on enzymatic activity. Soluble starch synthase III (Cre06.g282000; known as STA3 and re-named SSS3A in the *Chlamydomonas* v6.1 genome annotation), Starch Branching Enzyme 1 (Cre06.g289850; SBE1) and α-amylase 2 (Cre08.g362450; AMA2) were selected based on high peptide counts in the cGMP-functionalized beads. Additionally, *Chlamydomonas* mutants for STA3 (Ral et al., 2006, and references therein) and SBE1 (Tunçay et al., 2013) had been reported before, so that we hoped to be able to interconnect our biochemical with reported physiological data. In case of AMA2, we were intrigued by the predicted presence of an N-terminal ACT (<u>A</u>spartate kinase – <u>C</u>horismate mutase – <u>T</u>yrA) domain, as these domains are known to allosterically regulate enzymes upon the binding of small ligands (Grant, 2006; Lang et al., 2014).

### Recombinant STA3, SBE1 and AMA2 showed the expected catalytic properties

Our target proteins were heterologously produced in *E. coli* and purified by *Strep*-tag affinity chromatography. The activities of SBE1 and AMA2 were determined by analyzing their effects on starch- and, in case of SBE1, amylopectin-iodine complexes, whereas STA3 activity was analyzed by a coupled enzyme assay (Kulichikhin et al., 2016). During the first experiments we noted that all enzymes showed different specific activities depending on the amount of enzyme in the assay. Systematic testing of the optimal enzyme amounts revealed that SBE1 and STA3 were most efficient in the highest dilutions tested (0.5 µg enzyme per assay; Supporting Fig. S2). In case of AMA2, low amounts of enzyme (0.5 and 1 µg enzyme per assay) showed very high variability (Supporting Fig. S2A), which we assumed to be due to unpredictable protein instability. Based on these tests, we routinely employed 0.5 µg (3.6 pmol) of recombinant STA3, 0.5 µg (5.7 pmol) of SBE1 and 2 µg (25.3 pmol) of AMA2 in 400 µL reaction mixtures. All enzymes showed the expected catalytic activities (Fig. 1, Supporting Fig. S2) in that the addition AMA2 and SBE1 to starch or, in the case of SBE1, maize amylopectin, resulted in a decrease of the absorption of the corresponding carbohydrate-iodine complexes. AMA2 and SBE1 activities were tested on soluble potato starch and on storage starch isolated from N-deficient *Chlamydomonas* cells. On soluble potato starch, AMA2 exhibited a specific activity of 73.4 ± 1.4 U × µmol^-1^, and SBE1 of 467.1 ± 62 U × µmol^-1^, and both enzymes showed similar activities on *Chlamydomonas* starch (Fig. 1). On amylopectin, SBE1 developed an activity of 607.6 ± 149.4 U × µmol^-1^ (Fig. 1). STA3 activity in the presence of ADP-glucose and amylopectin resulted in the release of ADP as inferred from the generation of NADPH, and a specific activity of 529.3 ± 140.8 U × µmol^-1^ was determined (Fig. 1).

**Figure 1.**
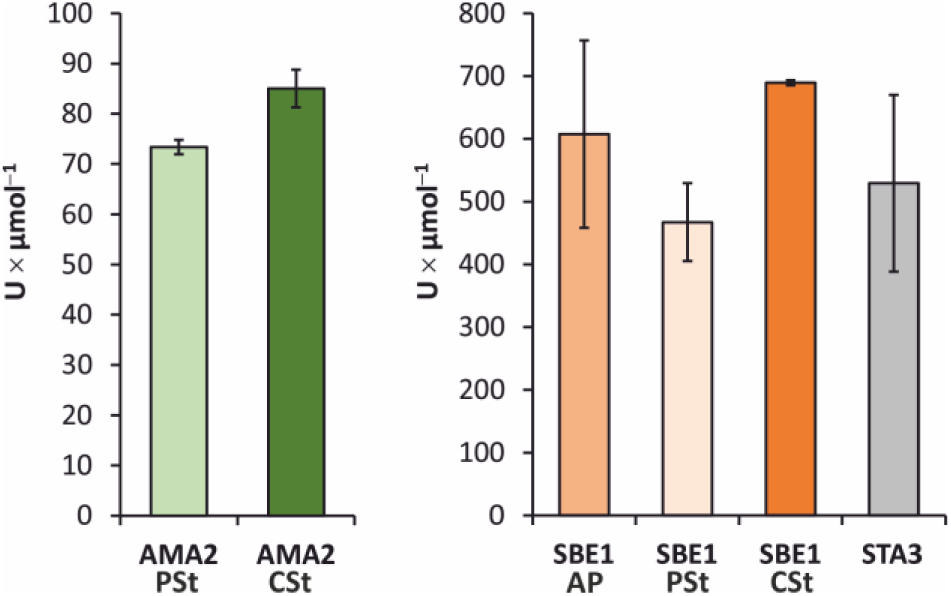
Recombinant AMA2, SBE1 and STA3 are active *in vitro*. *In vitro* activity assays were conducted employing 2 µg (25.3 pmol) AMA2, and 0.5 µg of SBE1 (5.7 pmol) or STA3 (3.6 pmol). Amylase and starch branching enzyme activities were determined on soluble potato starch (**PSt**), storage starch from N-deficient *Chlamydomonas* cells (**CSt**) or amylopectin (**AP**). AMA2 and SBE1 were diluted in 200 µL of ‘physiological buffer’ (50 mM potassium phosphate, pH 7.4, 50 mM KCl, 10 mM NaCl, 0.5 mM MgCl_2,_ 100 nM CaCl_2)_, mixed with starch or amylopectin as indicated beneath the x-axes, and incubated for 30 min at 30°C. The reactions were quenched by adding HCl, and the absorption was determined at λ = 580 nm (starch) or λ = 550 nm (amylopectin) after adding iodine solution. Enzyme activity (U) was defined as the decrease of 1 mg of iodine-stainable substrate per minute. STA3 activity was determined by an assay developed by Kulichikhin et al. (2016). In this case, U was defined as the release of 1 µmol of ADP per minute. The columns show average values, error bars the standard deviation. The assays were conducted with three independent enzyme preparations in three independent experiments, except for AMA2 and SBE1 in the presence of *Chlamydomonas* starch, for which the columns show means of two independent experiments, conducted with two independent enzyme preparations.

The specific activities of the enzymes were also analyzed in different pH values and under different temperatures to investigate whether they would be active within physiological ranges of these parameters. In case of AMA2 and SBE1, the pH optima lay in the neutral region with a tendency towards the alkaline (Supporting Fig. S3A, B). AMA2 activity was optimal at pH 8, but still high at pH 7.5 and pH 8.5 (Supporting Fig. S3A). On amylopectin, SBE1 exhibited nearly the same activities from pH 6.5 to pH 7.5, while its activities at pH 6 and pH 8 were still high, whereas it had a clearer optimum at pH 7.5 on starch (Supporting Fig. S3B). STA3 stood out in that its pH-dependent activity showed an optimum at pH 10. According to controls in which STA3 was replaced by ADP, the effect of the different buffer and pH values, respectively, employed for first reaction mixture, did only slightly affect the subsequent enzymes of the assay (Supporting Fig. S3C).

Temperature optima were determined for AMA2 and SBE1 and showed that both enzymes were active between 10°C and 50°C (Supporting Fig. S4). Similar to the pH-dependent activities, temperature-dependent activity assays resulted in a steeper graph in case of AMA2, with a temperature optimum at 40°C and a rather abrupt decrease of the specific activity at 45°C and 50°C (Supporting Fig. S4A). SBE1 showed a broader temperature profile and had the highest activities at temperatures of 30°C, 35°C and 40°C on amylopectin, and an optimum at 30°C on starch as substrate (Supporting Fig. S4B). On both substrates, SBE1 still showed rather high activities and 10°C and 45°C (Supporting Fig. S4B).

### AMA2 activity was sensitive to the thiol-oxidant diamide

Several enzymes of the starch metabolism exhibit different activities in the presence of thiol-modifying redox agents or thioredoxins (Kötting et al., 2010; Santelia et al., 2015; Skryhan et al., 2018). We tested the effect of the disulfide reductant DTT and the thiol-oxidizing agent diamide on the enzymes analyzed here (Fig. 2). The activities of SBE1 (Fig. 2A) and STA3 (Fig. 2B) did not change strongly after incubation with DTT or diamide. In contrast, AMA2 activity was notably impaired after incubation with diamide and decreased to 19.2 ± 9.2 % and 7.2 ± 2.7 % of the control activity after treatment with 1 and 5 mM diamide, respectively (Fig. 2C). In case of AMA2, we also tested whether its activity would be affected by the addition of calcium (Ca^2+^) or the chelator EDTA (Fig. 2D), because several α-amylases bind Ca^2+^ (e.g. Bush et al., 1989; Kadziola et al., 1994). The addition of Ca^2+^ to the standard activity assays did not result in enhanced AMA2 activities, but, in contrast, to moderately decreased activities that reached 86.1 ± 9.1 % of that of the control in the presence of 5 mM Ca^2+^ (Fig. 2D). The presence of EDTA did only have a strong effect at a concentration of 10 mM, resulting in an AMA2 activity of 64.0 ± 6.9 % relative to the control (Fig. 2D).

**Figure 2.**
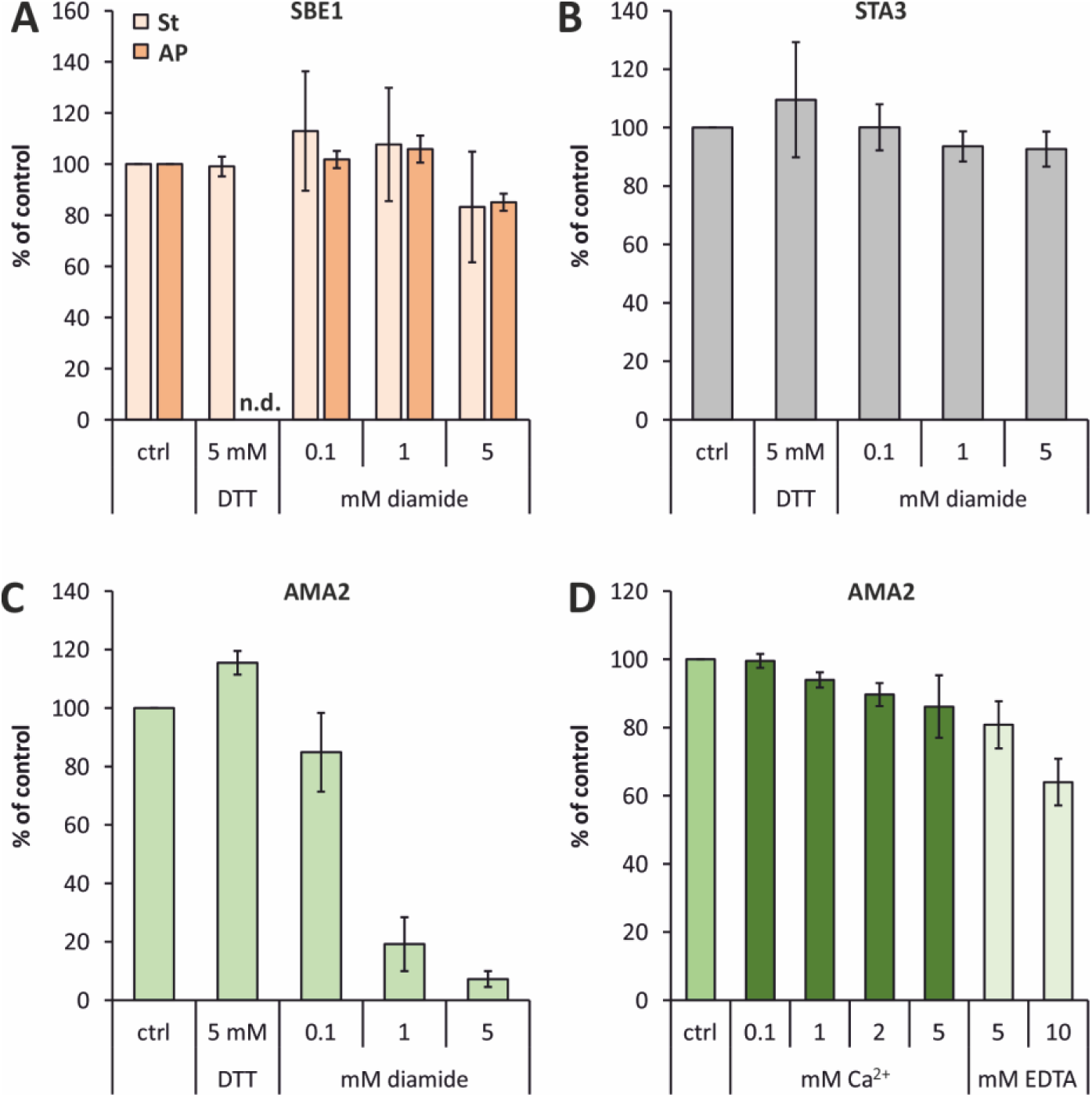
Effect of redox agents and calcium on enzyme activity. **A** to **C**: The enzymes were diluted to concentrations of 1 mg ×mL^-1 i^n 20 µL of ‘physiological buffer’ containing the indicated concentrations of dithiothreitol (DTT) or diamide or, as controls (**ctrl**), no supplements. After incubating the samples for 20 min at 23°C, aliquots containing 2 µg in case of AMA2 and 0.5 µg in case of SBE1 and STA3 were transferred to the standard activity assays that are described in detail in the caption of Fig. 1. SBE1 activity was tested on both substrates, starch (**St**) and amylopectin (**AP**). **D**: The effect of adding calcium (Ca^2+^) or ethylenediaminetetraacetic acid (**EDTA**) on AMA2 activity was tested by adding the indicated concentrations directly to the standard activity assay as described in the caption of Fig. 1. **A** to **D**: For each independent experiment, the mean of technical triplicates (duplicates in the case of the STA3 assay) of the control reactions was set to 100 %, and the activities determined in the presence of supplements were calculated accordingly. The columns show mean values of at least two independent experiments and at least two independent protein preparations. Error bars show the standard deviations. The specific activities of the controls were (**A**) 462.9 ± 83.7 U × µmol^-1 (^SBE1, St), 805.3 ± 29.9 U × µmol^-1 (^SBE1, AP), (**B**) 473.8 ± 26.8 U × µmol^-1 (^STA3), (**C**) 75.7 ± 10.5 U × µmol^-1 (^AMA2), and (**D**) 59.9 ± 15.7 U × µmol^-1 (^AMA2).

### None of the enzymes analyzed here was directly affected by (cyclic) nucleotides

Because we had assigned the starch metabolism enzymes analyzed here as possible cGMP binders based on our cGMP affinity chromatography approach, we tested whether cGMP would influence their activity. In neither case an effect of the presence of 100 µM cGMP on enzymatic activity could be observed (Supporting Fig. S5). We also tested cAMP, guanosine and adenosine monophosphate (GMP, AMP) as well as cyclic di-GMP (c-di-GMP) and guanosine tetraphosphate (ppGpp), because proteins regulated by cNMPs often show cross-reactivities such as in the case of cAMP- and cGMP-dependent protein kinases (Lorenz et al., 2017). The hyperphosphorylated guanosine nucleotide (p)ppGpp, originally described in prokaryotes, has a signaling function also in plants and algae (Field, 2018). Although a role of c-di-GMP in plants or algae has, to our knowledge, not been described, it regulates a glycogen-debranching-enzyme in *Streptomyces venezuelae* (Schumacher et al., 2022). However, none of the additional nucleotides or nucleotide derivates influenced the activities of AMA2, SBE1 or STA3 in the assays we employed during this work (Supporting Fig. S5). In case of STA3, a stimulating effect of AMP was very likely due to an impact on the subsequent enzymatic reactions, because it also resulted in a similarly enhanced NADPH generation when STA3 was replaced by ADP (Supporting Fig. S5C).

### AMA2 activity was enhanced in the presence of L-glutamine

Although we were unable to detect any direct effect on enzymatic activity by the tested nucleotides, we were still interested in the N-terminal ACT domain predicted for AMA2 because these domains are known for their interaction with small ligands (Grant, 2006; Lang et al., 2014). We first generated an AMA2 variant from which the N-terminal 269 amino acids were deleted, resulting in the α-amylase domain-only variant ΔAMA2 (Supporting Fig. S1). The presence of variant ΔAMA2 decreased the absorption of starch-iodine complexes, and, from the enzyme amounts tested, its optimal activity was reached when 0.5 µg (9.9 pmol) were employed (Supporting Fig. S2).Testing the activity of the ΔAMA2 variant at different pH values and temperatures showed that this α-amylase domain-only enzyme had similar pH and temperature optima as the full-length AMA2 enzyme (Fig. 3A, B), although its pH profile around neutral pH (pH 7 to pH 8.5) was broader (Fig. 3A). During these assays we noted that at physiological pH values (Fig. 3A) and at all temperatures tested (Fig. 3B), the ΔAMA2 variant exhibited a roughly 2.8-fold higher specific activity than AMA2.

**Figure 3.**
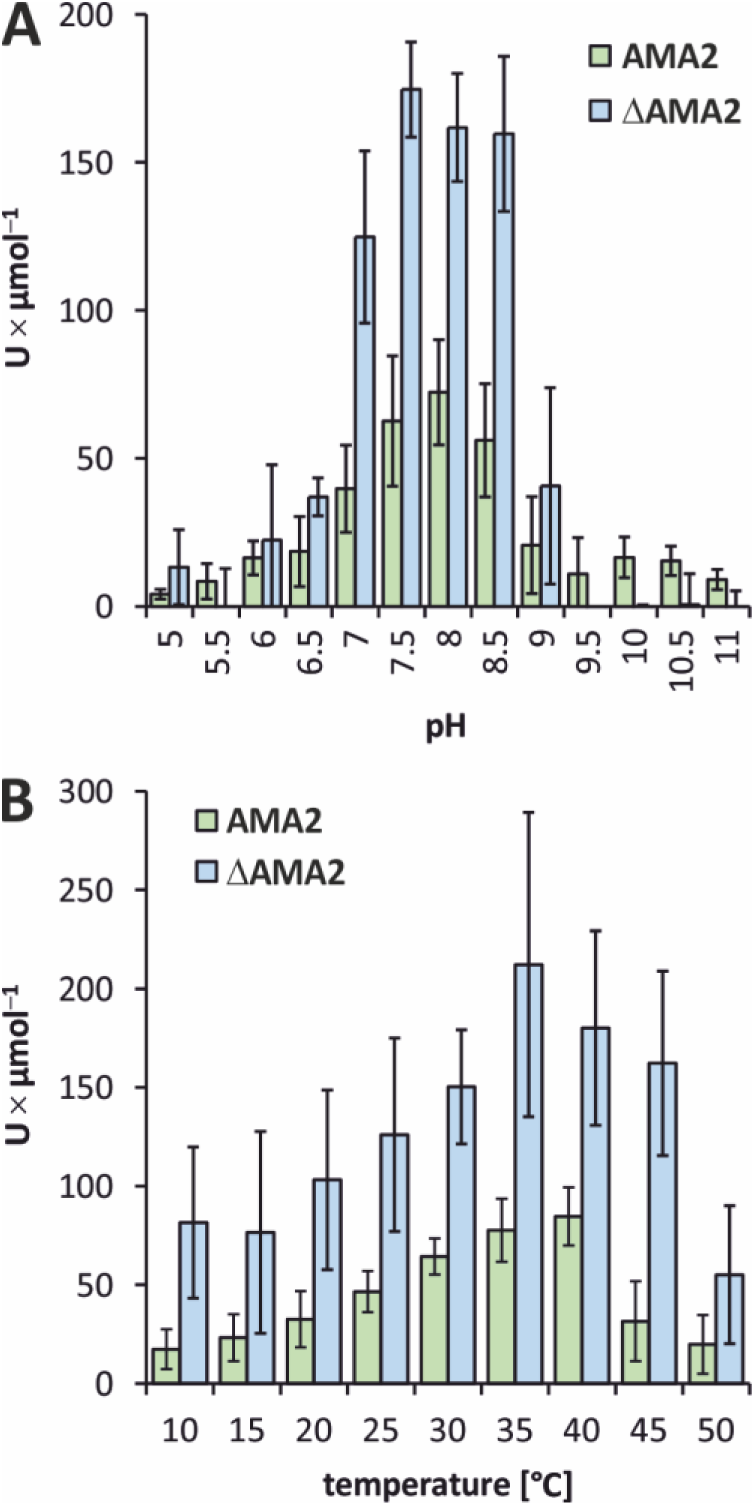
The α-amylase domain-only variant ΔAMA2 exhibits similar activity profiles as full-length AMA2. Amylase activities of AMA2 and the truncated variant ΔAMA2 were tested on soluble potato starch employing 2 µg (25.3 pmol) of AMA2 or 0.5 µg (9.9 pmol) of ΔAMA2. **A**: To determine pH optima, the proteins were added to 200 µl of Britton-Robinson buffer adjusted to the indicated pH values and mixed with 200 µL starch solution, corresponding to 140 µg of starch. After an incubation of 30 min at 30°C, the reaction was quenched by HCl, iodine solution was added, and the absorption was determined at λ = 580 nm. **B**: Temperature optima were determined as described for **A**, except that Britton-Robinson buffer, pH 7.5, was employed, and the incubation temperatures were adjusted as indicated at the x-axis. **A** and **B**: Columns show average values, and error bars indicate the standard deviations. AMA2 data are the same as shown in Supporting Figures S3 and S4. ΔAMA2 data were obtained with two independent enzyme preparations in three (**A**) or four (**B**) independent experiments.

ACT domains are often found in enzymes involved in purine or amino acid metabolism, and they allosterically regulate the enzymatic function by binding intermediates or end products of the respective pathway (Liberles et al., 2005). We tested the effect of all proteinogenic amino acids on the activities of AMA2 and the N-terminally truncated variant ΔAMA2 (Fig. 4; note that L-cysteine interfered with our assay and was therefore not included). The addition of L-arginine (Arg) and L-lysine (Lys) resulted in enhanced activities of both proteins to about 120 % (AMA2) and 140 % (ΔAMA2) (Fig. 4). The effect of the presence of L-glutamine (Gln) stood out in that it resulted in an activity of full-length AMA2 of 168.2 ± 2.4 % compared to the control reaction, while it hardly influenced the ACT domain-less ΔAMA2 variant whose activity reached 102.9 ± 6.8 % of that of its control activity (Fig. 4).

**Figure 4.**
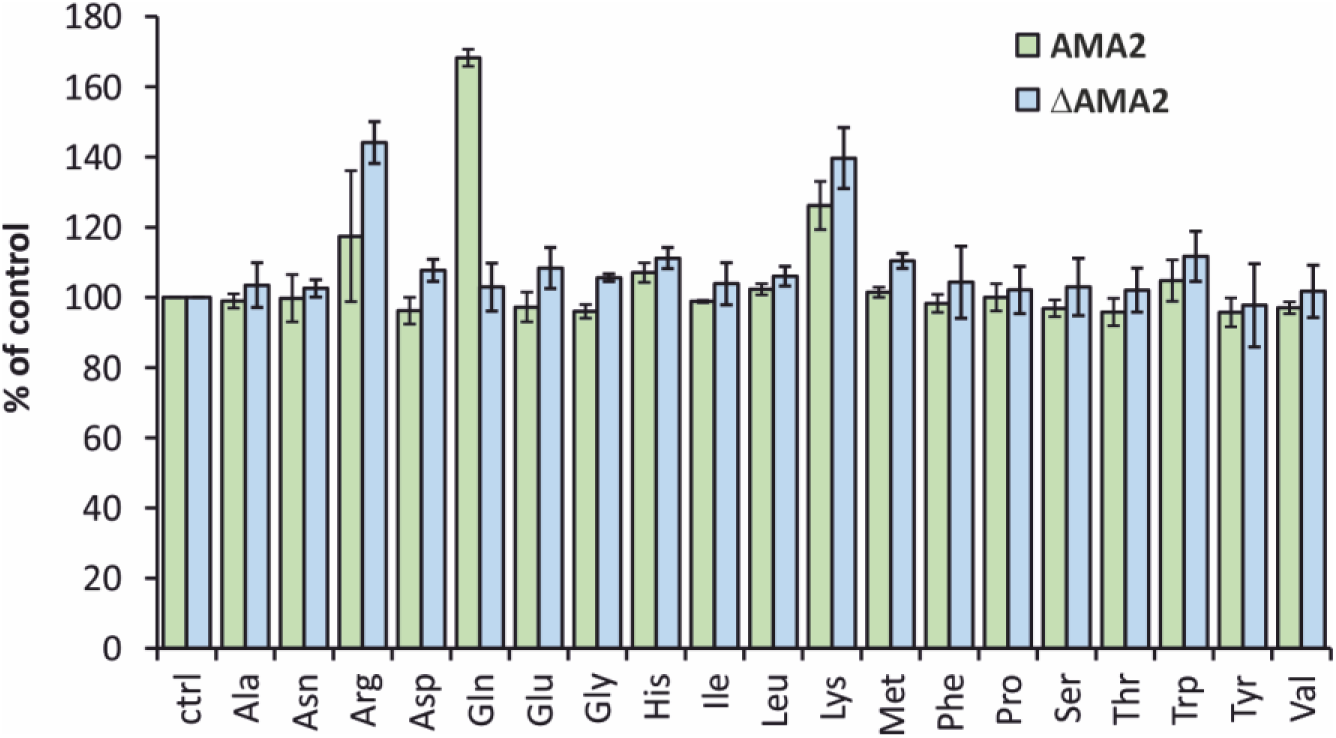
AMA2 activity is enhanced in the presence of L-glutamine. Activities of AMA2 (25.3 pmol) and the α-amylase domain-only variant ΔAMA2 (9.9 pmol) were determined as described in the caption of Fig. 1, except that the indicated L-amino acids were added to the reaction mixtures to a concentration of 5 mM. The averages of technical triplicates of the control reactions that did not include an amino acid (**ctrl**) were set to 100 % for each independent experiment, and the activities determined in the presence of individual amino acids were calculated in relation accordingly. Columns show mean values of at least two independent experiments, employing two independent protein preparations. Error bars show the standard deviations. The specific activities of the controls were 70.4 ± 4.5 U × µmol^-1 (^AMA2) and 179.3 ± 22.7 U × µmol^-1 (^ΔAMA2).

Testing the effect of the presence of different concentrations of Gln (0.05 to 10 mM) on AMA2 activity and calculating dose response curves resulted in a half maximal effective concentration (EC_50)_ of Gln of 2.1 ± 0.7 mM (Fig. 5A). Gln is a central intermediate and signaling molecule of the N assimilation pathway, and α-ketoglutarate (αKG) has been recognized as an important signaling molecule at the intersection of N and C metabolism (Huergo and Dixon, 2015). We therefore tested whether αKG influenced AMA2 activity, either alone or in combination with Gln. However, this was not the case (Fig. 5B).

**Figure 5.**
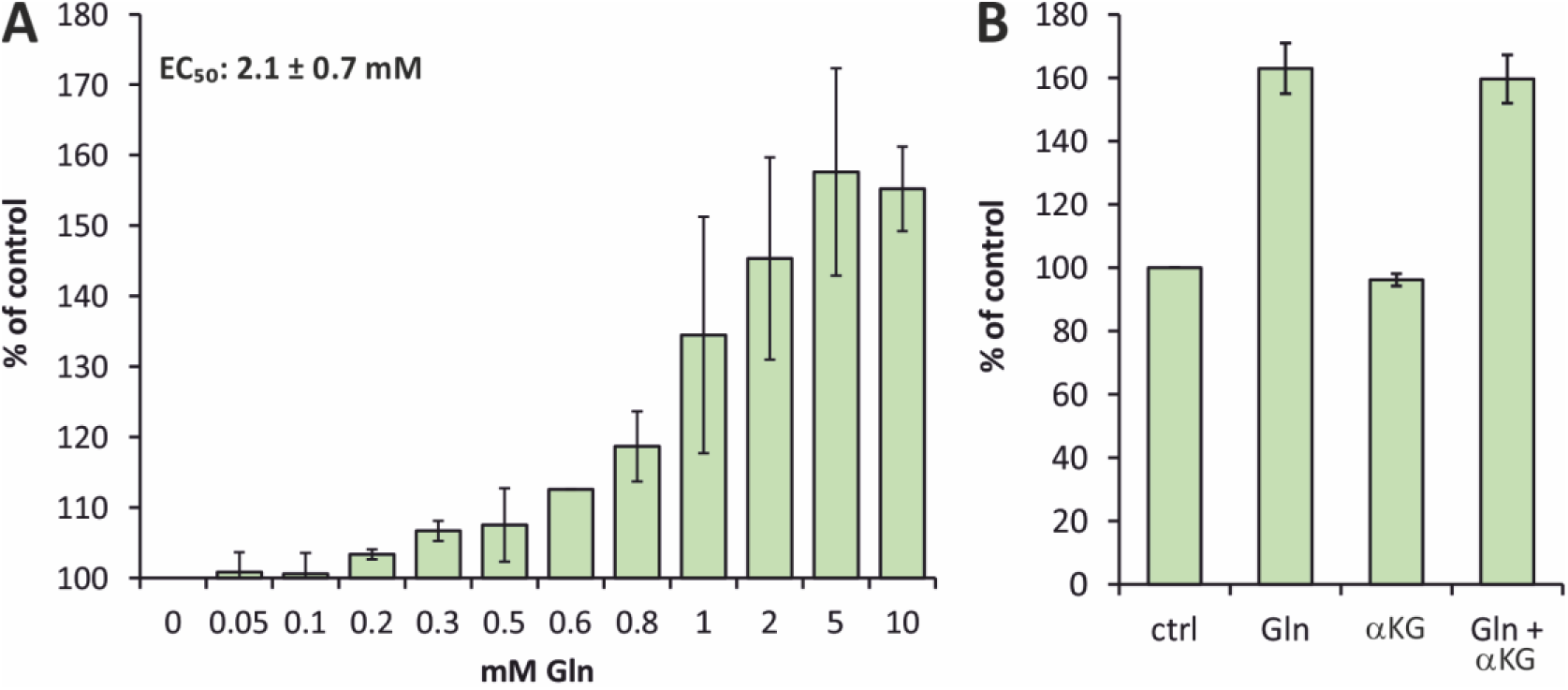
The effect of Gln on AMA2 activity is not altered in the presence of α-ketoglutarate. Activities of AMA2 (25.3 pmol) were analyzed as described in the caption of Fig. 4, except that different concentrations of L-glutamine (Gln) were added to the reaction mixtures (**A**) or that α-ketoglutarate (αKG) was added to a final concentration of 5 mM either alone or together with 5 mM of Gln (**B**). **A**: The effects of the indicated concentrations of Gln on AMA2 activity were tested in at least two independent experiments and at least two independent protein batches per concentration, but some concentrations were tested up to five times, using four enzyme preparations. A dose response curve was fitted employing the program OriginPro for each series independently, and the resulting values of the half maximal effective concentration (EC_50)_ were averaged. Note that the y-axis is scaled to begin at 100 %. **B**: The effect of the presence of Gln, αKG or both on AMA2 activity was tested at least thrice, employing at least three independent enzyme preparations. **A** and **B**: For each independent experiment, the specific activity of the control (**ctrl**), to which no supplement was added, was set to 100 %, and the specific activities of reactions in the presence of Gln or αKG were calculated accordingly. The specific activities of the controls were 61.6 ± 11.9 U × µmol^-1 (^**A**) and 67.4 ± 6.7 U × µmol^-1 (^**B**).

ACT domains often form inter- or intramolecular interactions (Lang et al., 2014). We applied size exclusion chromatography to test the oligomeric state of AMA2 in comparison to that of the N-terminally truncated variant ΔAMA2 (Fig. 6). Produced from the expression vector employed here, recombinant AMA2 and ΔAMA2 have calculated sizes of 79.2 kDa and 50.6 kDa, respectively. AMA2 eluted at a volume corresponding to a protein of 175 kDa, suggesting that it forms a dimer. According to its elution profile, the amylase domain-only variant ΔAMA2 was calculated to be a protein of 34 kDa, indicating that it was present as a monomer (Fig. 6). In both cases, small shoulders preceding the main elution fractions suggested the formation of a small fraction of tetramers (AMA2) or dimers (ΔAMA2). The binding of ligands usually does not result in different oligomeric states of ACT domain-containing proteins, but rather to conformational changes within the oligomer (Lang et al., 2014). We still tested whether the presence of a ten-fold molar excess of Gln would affect the elution profile of AMA2, however, this was not the case (Fig. 6).

**Figure 6.**
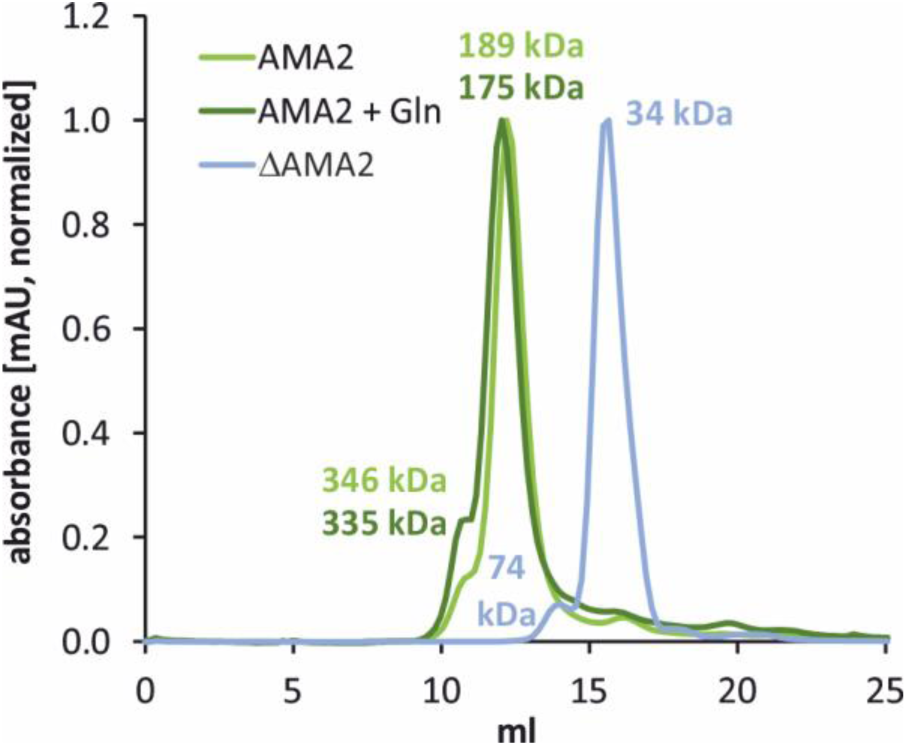
Recombinant AMA2 forms oligomers. Analytical size exclusion chromatography was performed to test for the oligomeric states of AMA2 and the α-amylase domain-only variant ΔAMA2. Proteins were diluted in Tris-HCl, pH 8, 150 mM NaCl, and loaded onto a Superdex 200 Increase 10/300 GL column with a 24-mL bed volume. Elution was done with the same buffer, and the absorbance of the eluate was recorded at λ = 280 nm. Normalized absorption profiles are shown. The molecular weights of the proteins present in the peaks were calculated according to a calibration performed employing a Gel Filtration Markers Kit. The elution profile of AMA2 was additionally tested in the presence of a ten-fold molar excess of L-glutamine (Gln). The molecular weights of the monomeric proteins as produced from the expression vectors are 79.2 kDa (AMA2) and 50.6 kDa (ΔAMA2).

### Selected amino acid exchanges in the AMA2 ACT domain affect amylase activity and Gln sensitivity

The ACT domain of AMA2 appears to mediate the activating effect of Gln (Fig. 4). We were interested to identify amino acids or parts of the ACT domain that would affect the Gln sensitivity of AMA2. The primary sequences of ACT domains are not well conserved (Aravind and Koonin, 1999; Liberles et al., 2005), which complicates the identification of key residues through comparisons with known ACT domains. To gain insights into how the ACT domain might fold, we modeled AMA2 as a dimer employing AlphaFold (note that an AlphaFold model of the AMA2 monomer is deposited in the AlphaFold Protein Structure Database under accession A0A2K3DGK5). Although the five models suggested by AlphaFold differed moderately, all showed the AMA2 dimer forming the shape of a fly, with the two ACT domains forming the ‘head’ and the amylase domains forming the ‘wings’ that are each connected by three α-helices to the ACT domains (Supporting Fig. S6A).

To analyze α-amylase- and ACT domain features in more detail, we employed the best-ranked AlphaFold model, and PyMOL for visualization. We aligned the AMA2 model with the crystal structures of Barley α-amylase 1 (abbreviated as HvAMY1 in the following; protein data bank (PDB) accession 1HT6) (Robert et al., 2003) as well as Barley α-amylase 2 in complex with the inhibitor acarbose (abbreviated as HvAMY2 in the following; PDB accession 1BG9) (Kadziola et al., 1998) (Supporting Fig. S6). Additionally, we aligned the primary sequences employing Clustal Omega (Supporting Fig. S7). The α-amylase domain of the AMA2 model superimposes both HvAMY1 and HvAMY2 structures well (Supporting Fig. S6A, B). For example, the residues of the catalytic Asp-Glu-Asp triad (Kuriki and Imanaka, 1999, and references therein) overlie almost completely (Supporting Fig. S6B, inset) and correspond to Asp476-Glu501-Asp584 of AMA2 (the numbering is according to the protein sequence Cre08.g362450.t1.2 including the N-terminal Met) (Supporting Fig. S7). Two of the three structural Ca^2+^-binding sites reported for both Barley α-amylases (Ca500 and Ca502 in PDB: 1HT6) (Robert et al., 2003, and references therein) appear conserved in the AMA2 model, although Lys435 would displace Ca500 (Supporting Fig. S6C). The residues that bind the third Ca^2+^, Ca501, are conserved neither in the structure model of AMA2 nor in its primary sequence (Supporting Fig. S6D, Supporting Fig. S7).

Because of the effect of diamide on AMA2 activity (Fig. 2C) we inspected its primary sequence and the AlphaFold model for Cys residues. The monomer contains nine Cys residues, two of which (Cys38, Cys45) are located within the putative chloroplast targeting peptide (Supporting Fig. S6A, Supporting Fig. S7). A third one (Cys97) is located before the second β-strand of the ACT domain (see below). Six Cys residues are located within the α-amylase domain, and Cys397 and Cys506, which lie near the active site, are visible on the surface of the AMA2 model (Supporting Fig. S6E).

The ACT domain of AMA2 is predicted by AlphaFold to encompass residues 86 to 171 and to follow a ββαββα topology (Supporting Fig. S8A). Archetypical ACT domains of enzymes involved in amino acid metabolism mostly follow a βαββαβ topology (Lang et al., 2014), but different arrangements of α-helices and β-strands have been described. The ββαββα topology is, for example, known from rat tyrosine hydroxylase (TyrH) (Zhang et al., 2014) or from ACT domains of several plant basic helix-loop-helix transcription factors (Feller et al., 2017; Lee et al., 2023). The typical arrangement of four antiparallel β-strands and two antiparallel α-helices to one side of the β-sheet, however, is mostly conserved, and also predicted for the AMA2 ACT domain (Supporting Fig. S8A).

Structures of ACT domain-containing proteins show that ACT domains mediate a multitude of oligomerizations (Grant, 2006; Lang et al., 2014). Often, ligands bind at interfaces of ACT domains, but they can also be coordinated by residues within a single ACT domain. One example for both modes is *Arabidopsis* aspartate kinase, in which one effector, Lys, binds at the ACT domain interface, and the other, SAM, binds within the ACT domain (Mas-Droux et al., 2006). Alignment methods suitable for the diverse primary sequences of ACT domains revealed conserved glycine (Gly) residues in loops between regions with secondary structures that, in some cases, were demonstrated to be involved in ligand binding (Grant, 2006, and references therein). A Gly-to-aspartate (Asp) exchange in *Arabidopsis* serine/threonine/tyrosine (STY) kinase (STY8), for example, which is regulated by both isoleucine (Ile) and SAM, results in a loss of sensitivity to Ile (Eisa et al., 2019). Inspecting the structure model of the AMA2 ACT domain, we identified three Gly residues in loops, namely G122 (between α-helix 1 and β-strand 3), G135 (between β-strands 3 and 4) and G148 (between β-strand 4 and α-helix 2) (Supporting Fig. S8B). AMA2 variants were generated in which these Gly residues were exchanged by Asp, and their activity was determined as described above for the wild type enzyme both in the absence and presence of Gln. All variants had nearly the same specific activities as the wild type protein, and their activities were increased in buffer containing Gln (Fig. 7A). Variants G122D and G148D showed a similar response to Gln as the wild type enzyme, while the increase of activity of variant G135D was stronger, reaching 210 ± 11 % of its activity in the absence of Gln (Fig. 7A).

**Figure 7.**
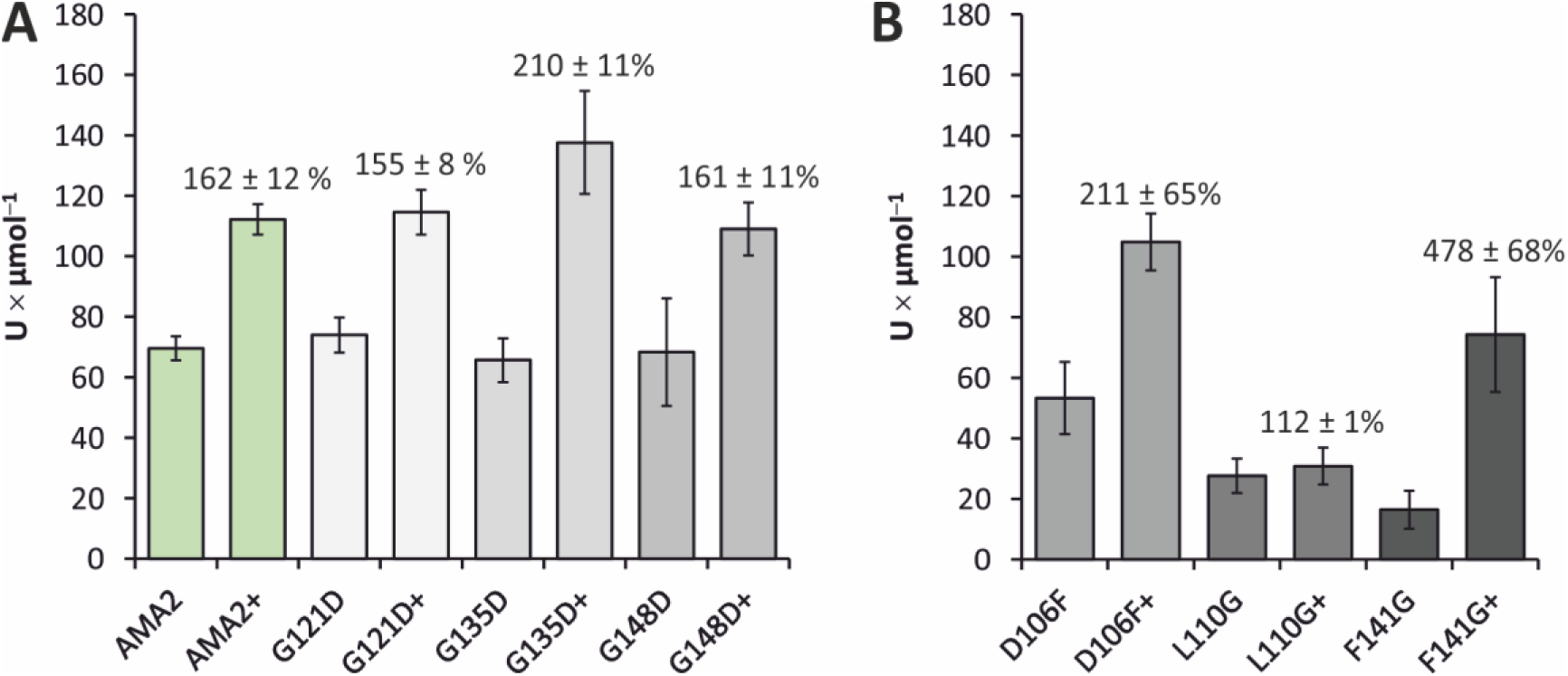
Amino acid exchanges in the AMA2 ACT domain have different effects on activity and Gln sensitivity. Activities of the AMA2 wild type enzyme and single amino acid exchange variants were determined as described in the caption of Fig. 4, both in the absence and presence of 5 mM of Gln (the latter is indicated by a plus sign after the variant denomination). Each enzyme variant was tested in at least two biological replicates and two independent experiments. Columns show the averages of the specific activities, error bars indicate the standard deviation. For each variant, the relative activity determined in the presence of Gln compared to that in the absence of Gln was calculated and is indicated above the columns. **A**: Activities of AMA2 variants in which glycine (G) residues within loops of the ACT domain were exchanged (see Supporting Fig. S8B). **B**: Activities of variants in which conserved residues identified by an alignment (Supporting Fig. S9) were exchanged (depicted in Supporting Fig. S8C).

We then decided to search for ACT domains with sequence homology to the AMA2 ACT domain to possibly identify conserved residues that might be important for its function. As described in detail in the materials and methods section and in the captions of Supporting Table S1 and Supporting Fig. S9, we retrieved similar sequences employing NCBI’s BlastP tool. Before generating the alignment, we manually added the ACT domain sequences of three proteins: *E. coli* GlnD is a uridylyltransferase/uridylyl-removing enzyme that modifies the P_II p_rotein, a central signal transduction protein (Forchhammer et al., 2022), in response to Gln, for which the ACT domains of GlnD are required (Zhang et al., 2010). ACR11 is one of twelve so-termed ACT repeat (ACR) proteins in *Arabidopsis* and is involved in the regulation of the GS/GOGAT cycle that assimilates ammonium and forms Gln (Liao et al., 2020, and references therein). We also added the ACT domain of *Chlamydomonas* starch phosphorylase STA4 (PhoB), which was also present in our cGMP affinity chromatography selection (Supporting Data Set 1, Sheet 4).

Inspecting the alignment but focusing on the ACT domain sequences of *E. coli* GlnD and *Arabidopsis* ACR11, we selected three additional residues that we exchanged in AMA2, namely Asp106, Leu110 and Phe141 (the numbering is according to full-length AMA2). Residue Asp106 was found in both ACT domains each of *E. coli* GlnD and *Arabidopsis* ACR11 and is represented by Asp or Asn throughout our alignment (Supporting Fig. S9). Leu110 is also quite conserved, being present in most of the sequences, whereas Phe141 is strictly conserved (Supporting Fig. S9). According to our AlphaFold model, Asp106 and Leu110 are located at loop regions, whereas Phe142 reaches in the space between β-strand 4 and α-helix 1 (Supporting Fig. S8C). Although the hydrophobic side chain of Phe is not intuitively associated with coordinating a polar amino acid, the aliphatic part of Gln is sandwiched between two Phe residues in a Gln-binding protein (Sun et al., 1998). The activity of the AMA2 variant D106F, in which Asp106 was exchanged by Phe, was very similar to that of the wild type enzyme and it was still sensitive towards Gln (Fig. 7B). The enzyme variants in which Leu110 and Phe141 were individually exchanged by Gly (L110G; F141G) showed impaired activities and contrary changes of activity in the presence of Gln: AMA2 L110G was almost insensitive to Gln, reaching 112 ± 1 % of its activity determined in the absence of Gln, whereas the F141G variant was stimulated to 478 ± 68 % of its standard activity when Gln was present in the assay (Fig. 7B).

## Discussion

### The enzymes analyzed here have mostly typical biochemical characteristics

The three enzymes that we picked from our cGMP affinity chromatography were detected in the *Chlamydomonas* chloroplast (Terashima et al., 2010), suggesting that they are involved in the plastid starch metabolism of *Chlamydomonas*, and all showed the expected activities in their recombinant forms (Fig. 1). The α-amylase AMA2 and the starch-branching enzyme SBE1 catalyzed changes to the starch- or, in the case of SBE1, amylopectin substrate as indicated by the absorbance of polyglucan-iodine complexes. SBE1 was characterized in its recombinant form before and shown to be active on amylose (Courseaux et al., 2023). Both enzymes were also active on native *Chlamydomonas* storage starch, suggesting that they would be active *in vivo* on insoluble starch granules. The soluble starch synthase STA3 was capable of utilizing ADP-glucose as determined by the conversion of ADP to NADPH by subsequent enzymatic reactions in an assay developed previously (Kulichikhin et al., 2016). In the case of SBE1 and STA3, we noticed clear trends towards higher efficiencies when the enzymes were more dilute (Supporting Fig. S2), which might simply be due to a better access of individual protein molecules to the polyglucan substrates. AMA2, however, revealed high standard deviations of its specific activities in higher dilutions (Supporting Fig. S2). This suggests that the enzyme was instable under these conditions. As we found recombinant AMA2 to be present as a dimer (see below; Fig. 6), one possibility is that the complex dissociated at lower protein concentrations. In view of this behavior, we chose to employ higher concentrations of AMA2 in all activity assays, so that direct comparison between full-length AMA2 and its N-terminally truncated variant ΔAMA2 (see below) need to be interpreted with caution.

AMA2 and SBE1 showed pH optima in the neutral region (Supporting Fig. S3), although AMA2 revealed a steeper profile with a clear optimum at pH 8. This suggests that, in living *Chlamydomonas* cells, AMA2 would be more active upon illumination when photosynthetic electron transport results in an alkalization of the chloroplast stroma (also see below). STA3 showed a broad pH profile with an optimum at pH 10 but still high activities at both pH 5 and pH 12 (Supporting Fig. S3). High activities of soluble starch synthases in the alkaline region were noted before (e.g. Imparl-Radosevich et al., 1999). Although intracellular pH values are commonly in the neutral region, it is well possible that the microenvironment of starch synthases requires adaptability to different proton concentrations. In case of *Chlamydomonas*, starch metabolism is additionally associated with the pyrenoid, a phase-separated compartment within the chloroplast responsible for CO_2 c_oncentration. The pyrenoid is surrounded by starch sheaths, and proteomics have identified many starch metabolism enzymes in this subcompartment (Mackinder et al., 2017; Zhan et al., 2018; Lau et al., 2023) (also see below). Indeed, SBE1 was present in the pyrenoid proteome (Zhan et al., 2018), and STA3 in the pyrenoid ‘proxisome’ (Lau et al., 2023). The crowded environment of the liquid-like organelle might also result in (locally) different pH values.

### Starch metabolism enzymes might indirectly bind to cGMP

The activity of none of the enzymes we characterized here was influenced by the presence of cGMP or the other nucleotides tested (Supporting Fig. S5), although the proteins were retained with high peptide counts in cGMP-functionalized agarose beads. This suggests that either these enzymes bound unspecifically or that they were present because of interactions with other proteins. Principally, the approach was suitable to capture proteins that bind cNMPs. Of the 66 proteins we assigned as possible (indirect) cGMP binders (Supporting Data Set 1, Data Sheet 4), four are predicted to be typical cNMP-binding proteins in that they contain CNB or GAF domains (Aravind and Ponting, 1997; Rehmann et al., 2007; Heikaus et al., 2009). These proteins were only detected in the 2-AH-cGMP- and not in the 2’-AHC-cGMP bead fractions, suggesting that the former, to which the cGMP moiety is tethered through its C2 amine group, were better suited to capture canonical cNMP-binding proteins. Typically, cNMPs are mostly buried within GAF domains (Heikaus et al., 2009), and in mouse PDE2a, for example, only the C2 amino group of the guanosine base protrudes to the protein surface (Martinez et al., 2002) (Supporting Fig. S10A). A larger part of cNMPs bound to CNB domains can be accessible from the protein surface, however, the ribose moiety projects towards the protein interior (e.g. Kim et al., 2016) (Supporting Fig. S10B). This might hinder an efficient binding to the 2’-AHC-cGMP beads, in which the tether to the agarose material is attached at the 2’ C-atom of the ribose moiety.

Of the remaining possible cGMP binders, many proteins are known or predicted to bind nucleotides, nucleotide-derivatives or -intermediates, or nucleic acids. Therefore, the proteins involved in starch metabolism captured by the cGMP-functionalized beads might have bound unspecifically. As noted in the introduction, starch metabolism enzymes can form multi-protein complexes (Kötting et al., 2010; Geigenberger, 2011; Schreier et al., 2019; Abt and Zeeman, 2020) so that it is possible that the proteins captured were bound to the beads through ADP-binding soluble starch synthases, or an ATP-binding protein such as the alpha-glucan, water dikinase GWD2 (Supporting Data Set 1, Data Sheet 4). *Chlamydomonas* starch-branching enzymes were indeed detected in high-molecular weight complexes (Courseaux et al., 2023). In the case of SBE1 and STA3, possible protein partners were identified through high-throughput affinity purification approaches: SBE1 was affinity-purified by Rubisco small subunit (RBCS) 2, and STA3 by starch branching enzyme 3 by Mackinder et al. (2017), and Wang et al. (2023) found SBE1 interacting with RBCS1.

Alternatively, the starch metabolism enzymes detected in our cGMP interactome might have bound to the cGMP beads through an interaction with a cNMP-dependent protein kinase. We detected phosphorylated peptides of the three enzymes selected here in *Chlamydomonas* phosphoproteome studies (AMA2 and SBE1 in Shinkawa et al. (2019), and AMA2 and STA3 in Wang et al. (2014)), demonstrating that they are phosphorylated *in vivo*. The *Chlamydomonas* genome encodes for several protein kinases with predicted CNB domains, three of which we detected in the 2-AH-cGMP beads. FAP19 (Cre02.g076900) is involved in flagellar signaling (Wang et al., 2006), whereas FAP358 (Cre12.g493250) and FAP295 (Cre16.g663200) have, to our knowledge, not been analyzed yet. None of these three proteins is predicted to have a chloroplast targeting sequence. Notably, however, FAP358 and FAP295 were detected by the affinity purification approach of Mackinder et al. (2017) mentioned above. FAP358 was affinity-purified by the bicarbonate transporter HLA3, the putatively chloroplastic acyl-carrier protein ACP2 and the thylakoid membrane bestrophin Cre16.g662600, although the scores (all < 2.33) were much lower as the cut-off set by the authors for high-confidence interactions (> 47.516). FAP295 was pulled down by NAR.2/LCIA, an anion transporter at the chloroplast membrane (score 32.76), also by HLA3 (score 28.89), by the CO_2 c_hannel LCI1 (score 4.67) and also by bestrophin Cre16.g662600 (score 3.3). As HLA3 and LCI1 are plasma membrane transporters, both protein kinases were affinity-purified by both chloroplastic and plasma membrane proteins. Thus, although we find it intriguing to speculate on an interaction of cNMP-dependent protein kinases with starch metabolism enzymes, this idea will have to be tested in the future.

### AMA2 activity is enhanced by L-Gln through its ACT domain

When we started experiments on the selected starch metabolism enzymes, the N-terminal ACT domain of AMA2 was our first candidate for a possible interaction with cGMP. However, as none of the nucleotides tested here had an effect on AMA2 activity, we tested for an influence of proteinogenic amino acids, because these are common ligands of ACT domains (Liberles et al., 2005; Lang et al., 2014; Eisa et al., 2019). The α-amylase domain-only variant ΔAMA2 was tested in parallel to determine effects likely to exert influence through the ACT domain. Arg and Lys stimulated the activity of both proteins (Fig. 4) so that these two amino acids probably had unspecific effects, for example by enhancing the solubility of the proteins, although an effect on the α-amylase domain cannot be excluded. Gln, in contrast, only stimulated the activity of the full-length enzyme (Fig. 4), suggesting that it exerts its effect through the ACT domain. In our assays, the variant ΔAMA2 showed higher specific activities (Supporting Fig. S2, Fig. 3) so that it may be hypothesized that the ACT domain has an inhibitory effect on α-amylase activity, which is released upon Gln-binding. However, as discussed above, we assayed different protein amounts of AMA2 and ΔAMA2. Additionally, it is possible that the smaller ΔAMA2 variant simply had a better access to the starch substrate than the larger and dimeric (see below) AMA2 protein.

Both the AMA2 model in the AlphaFold Protein Structure Database (A0A2K3DGK5) as well as our AlphaFold model of an AMA2 dimer predict a typical ACT domain fold of the N-terminal region (Supporting Figs. S6 and S8). Our gel filtration analyses indicate that full-length AMA2 forms at least a dimer, both in the absence and the presence of Gln, while the ΔAMA2 variant eluted as a monomer (Fig. 6), suggesting that the ACT domain is involved in the oligomerization of AMA2. Notably, AMA2 was found to affinity-purify itself in the above-mentioned immunoprecipitation approach of Wang et al. (2023), corroborating our SEC elution profile. It is common that ACT domains mediate protein-protein interactions, and that ligand-binding changes the conformation of, but does not disintegrate, the oligomer (Grant, 2006; Lang et al., 2014). Also, the formation of an eight-stranded β-sheet by the side-by-side arrangement of two ACT domains as predicted by our dimeric model was already observed in the first crystal structure of an ACT domain-containing protein (Schuller et al., 1995) and often since (Grant, 2006; Lang et al., 2014). Notably, however, in many cases the α-helices face outwards, which contrasts with our model (Supporting Fig. S6). Whether this is the true AMA2 structure and, if so, whether it has a specific functional consequence will require future studies.

To identify the Gln-binding site within the AMA2 ACT domain we generated six single amino-acid exchange variants based on comparisons to known and predicted ACT domains, the latter identified by homology searches (Supporting Table S1, Supporting Fig. S9). Neither AMA2 activity nor the stimulating effect of Gln were strongly affected by the exchanges of three Gly residues in loops (Fig. 7A; Supporting Fig. S8B, C), suggesting that these residues are vital neither for the enzyme’s active structure nor for the impact of Gln. However, the specific activities of variants with the exchanges D106F and particularly L110G and F141G were lower than that of the wild type enzyme (Fig. 7B). Additionally, the impact of Gln on the latter two variants varied strongly from that on the wild type protein, albeit to opposite effects: The activity of variant L110G was rather insensitive to the presence of Gln, while that of variant F141G increased more than four-fold (Fig. 7B). We hesitate to assign a Gln-coordinating function to L110G, because the impaired activity of this variant argues for a structural disturbance. However, all the latter three residues as well as G135, whose exchange resulted in a slightly stronger stimulating effect of Gln (Fig. 7A) are located in or around a wedge-like shape formed between β-strands 3 and -4 and α-helix 1 (Supporting Fig. S8A, C), suggesting that this is the region were Gln binds.

### Integration of starch degradation and Gln levels – a day in the life of *Chlamydomonas* R

The activating effect Gln exerts on AMA2 activity suggests that the α-amylase AMA2 might be a direct, enzymatic means to coordinate C- and N-metabolism in *Chlamydomonas*. In *Chlamydomonas* cells grown in day/night cycles, the starch content shows a rhythmic pattern. Depending on growth conditions (autotrophic or photomixotrophic), starch levels are highest at the end of the subjective day or a few hours after the onset of darkness, and decrease thereafter (Klein, 1987; Thyssen et al., 2001; Ral et al., 2006). In both autotrophic and mixotrophic growth conditions, net starch degradation usually continues well into the next light period (Klein, 1987; Thyssen et al., 2001; Ral et al., 2006; Strenkert et al., 2019), which was suggested to corelate with the cell cycle (Ral et al., 2006). *Chlamydomonas* cultures grown in regular dark/light cycles synchronize their cell-cycle, exhibiting cell growth during the day and cell division in the beginning of the night (Bišová and Zachleder, 2014; Cross and Umen, 2015). Accordingly, cellular protein and chlorophyll contents increase during the day (Thyssen et al., 2001; Strenkert et al., 2019). RNA also accumulates rather constantly during the light phase, while the cellular DNA content increases towards the evening (Grant et al., 1978; Jüppner et al., 2017; Pokora et al., 2017). Nitrogen is present in considerable amounts in all of these molecules, and nitrate assimilation as well as ammonium fixation enzymes are indeed highly active during the day, although with different activity profiles: Both nitrate- and nitrite reductases (NR and NiR, respectively) activities peak during the light phase, but whereas NR is inactive during the night, NiR maintains a constant activity (Martínez-Rivas et al., 1991). The activities of glutamine synthetase (GS) and ferredoxin-dependent glutamine oxoglutarate aminotransferase (GOGAT, or glutamate synthase) increase constantly during the day and stay on a high level in the night (Martínez-Rivas et al., 1991). Glutamine, glutamate and 2-oxoglutarate levels are higher during the light period, all three showing two maxima during the subjective day, but stay on relatively high levels during the night (Jüppner et al., 2017).

In view of these data, it seems reasonable to suggest that, during the day, *Chlamydomonas* coordinates N- and C assimilation to efficiently biosynthesize N-containing molecules required for optimal cell growth. AMA2 could be one of the coordinating hubs, in that available N in the form of Gln stimulates its activity to release C-skeletons. Our observations that recombinant AMA2 shows highest activity at pH 8 (Supporting Fig. S3) and is nearly inactive after pre-treatment with the thiol-oxidizing agent diamide (Fig. 2) support the notion that it might be mostly active during the day, similar to what has been observed for *Arabidopsis* α-amylase AMY3 and β-amylase BAM1 (Skryhan et al., 2018, and references therein). Reversible Cys reduction is employed by photosynthetic organisms to convey active photosynthetic electron transport to downstream sinks such as the CBB cycle, often by the thioredoxin system (Michelet et al., 2013). AMA2 contains several Cys residues (Supporting Fig. S7) and was identified in a ‘thioredoxome’ study (Pérez-Pérez et al. (2017); AMA2 is listed by its previous UniProt identifyer A8J4D3), implying that the enzyme might be regulated through the thioredoxin system. Two of the Cys residues of AMA2 are located near the active site and appear solvent accessible as judged by viewing the surface of the AlphaFold AMA2 dimer model (Supporting Fig. S6E). In the model, none of the Cys residues appear close enough to allow the formation of intra- or intermolecular disulfides, however, it is possible that the protein forms higher-order oligomers under oxidizing conditions, or that only single Cys thiols are modified. AMA2 was indeed also identified by proteomics studies that captured *S*-glutathionylated (Zaffagnini et al., 2012) or *S*-nitrosylated proteins (Morisse et al., 2014) (in both studies, AMA2 can be found under its RefSeq identifier XP_001696014). It should be noted that the preceding argumentation is based on the hypothesis that AMA2 is located in the plastid. As mentioned above, AMA2 was detected in the chloroplast by proteomics (Terashima et al., 2010), and the localization prediction tool PredAlgo (Tardif et al., 2012) predicts a chloroplast targeting peptide (also see Supporting Fig. S1). However, a recent high-throughput approach that employed fluorescent protein-tagging detected AMA2 at the nuclear envelope (Wang et al., 2023). Although the authors discussed the possibility of mis-localizations, future studies of AMA2 need to pinpoint its localization(s).

## Conclusion

Our biochemical data indicate that the ACT domain of AMA2 functions as an amino acid sensor as has been demonstrated for many ACT domains before. We identified several putative α-amylases with N-terminal ACT domains in algae (Supporting Table S1), suggesting that the co-occurrence of these two domains is widespread in algae. We also detected a predicted N-terminal ACT domain in the *Chlamydomonas* starch phosphorylase STA4 (PhoB), although its effect, if any, on phosphorylase activity has, to our knowledge, not been analyzed. STA4 appears to be solely involved in the accumulation of storage starch (Dauvillée et al., 2006), so that its ACT domain might fulfil another role as that in AMA2, which we hypothesize to be involved in starch degradation.

Gln as a central indicator for the N status also binds to the *Chlamydomonas* P_II p_rotein and thereby indirectly relieves *N*-acetyl-L-glutamate kinase (NAGK), which is central for polyamine and ornithine biosynthesis, from Arg feedback inhibition (Chellamuthu et al., 2014). Notably, the EC_50 v_alue of 2.1 ± 0.7 mM of Gln we determined for AMA2 activation (Fig. 5A) is close to the EC_50 o_f Gln Chellamuthu et al. (2014) determined for the activation of NAGK by the P_II p_rotein (2.4 ± 0.8 mM), suggesting that both regulatory effects might be operative under similar physiological conditions. Rather recently, it was shown that *Chlamydomonas* phosphoenolpyruvate carboxykinase isoform 2 is inhibited by Gln (Torresi et al., 2023), representing an additional example of a direct effect of Gln. It appears that the alga employs Gln as a signaling molecule to coordinate N- and C metabolism directly at the enzymatic level in diverse contexts, and that AMA2 might be an additional example.

## Supporting information

Supporting Data Set 1

## Acknowledgments

We thank the Deutsche Forschungsgemeinschaft (DFG; German Research Foundation) for funding within RTG 2341 (Microbial Substrate Conversion (MiCon); A.H., E.H.). A.P. acknowledges funding from the German Federal Ministry of Education and Research, within the ERA-MIN2 framework, project MiCCuR (033RU011B). We are very thankful for excellent technical assistance and advise during the cGMP affinity chromatography experiments from M.Sc. Sabrina Duda (Photobiotechnology group, Ruhr University Bochum), Dr. Jan Lambertz (Department of Plant Biochemistry, Ruhr University Bochum) and Prof. Dr. Marc Nowaczyk (currently at the Department of Biochemistry, University of Rostock, Germany).

## Author contribution

L.S. and A.H. designed the research; L.S. performed the research; all authors analyzed data; L.S. and A.H. wrote the first draft of the manuscript; all authors edited the manuscript.

## Conflict of interest

The authors declare no conflict of interest.

## Supporting Information

**Supporting Table S1.**
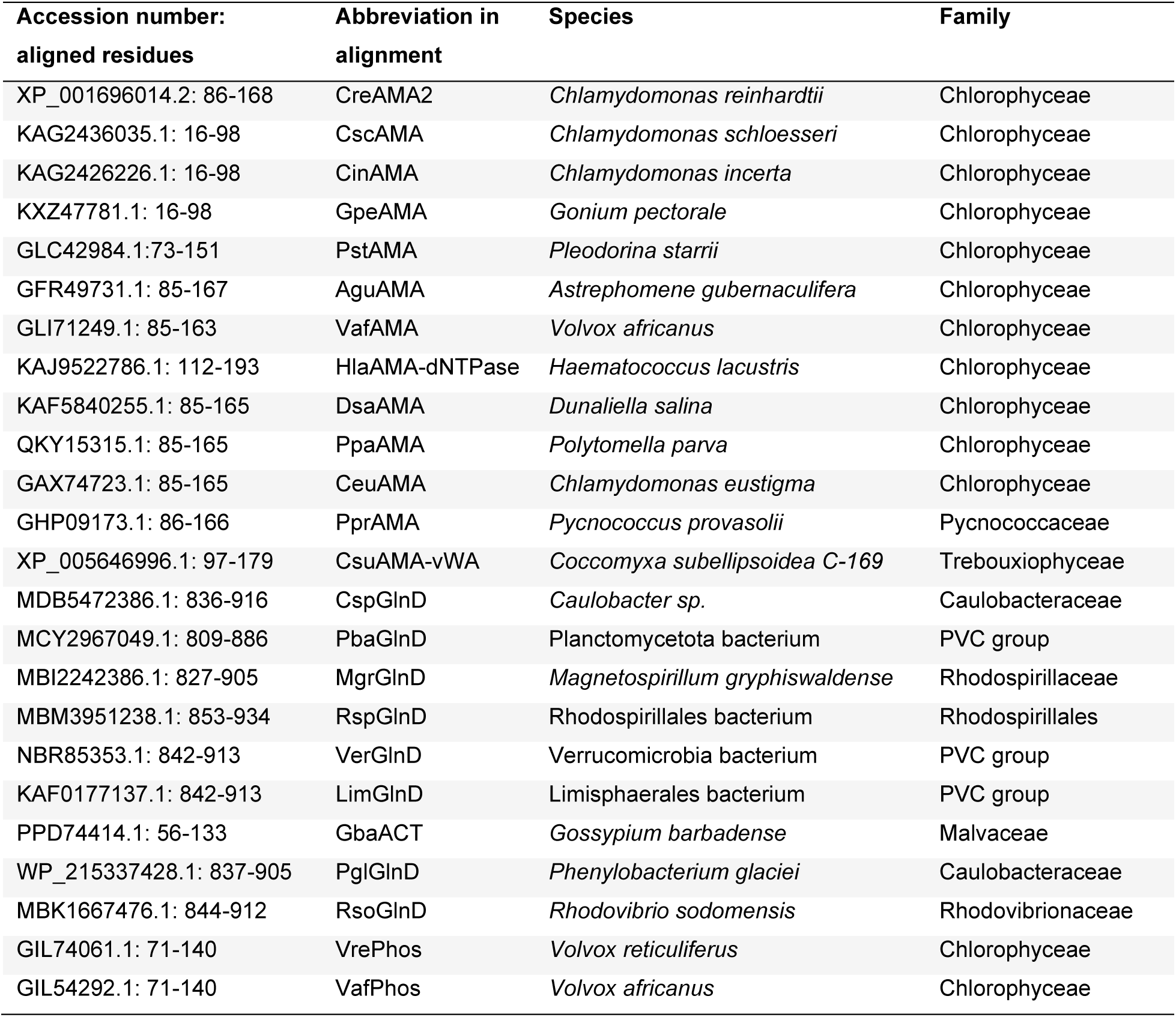
Protein sequences used for the alignment of ACT domains. The primary sequence of the AMA2 ACT domain was employed to retrieve similar sequences by NCBI’s BlastP tool, and manually selected sequences were used to generate an alignment by Clustal Omega (Supporting Fig. S9) (see the materials and methods section for details). NCBI accession numbers and aligned residues of the protein sequences employed for the alignment are indicated in the first column. Full-length primary sequences of the selected ACT domain proteins were analyzed for additional domains, which are indicated as extensions of the species abbreviations as shown in the second column and abbreviated as follows: **AMA**: alpha amylase, **dNTPase**: p-loop containing nucleoside triphosphate hydrolase, **vWF**: von Willebrand factor type A domain, **GlnD**: [protein-PII] uridylyltransferase/uridylyl-removing enzyme, **ACT**: ACT domain-only (plant ACR proteins), **Phos**: starch / glycogen phosphorylase.

**Supporting Figure S1.**
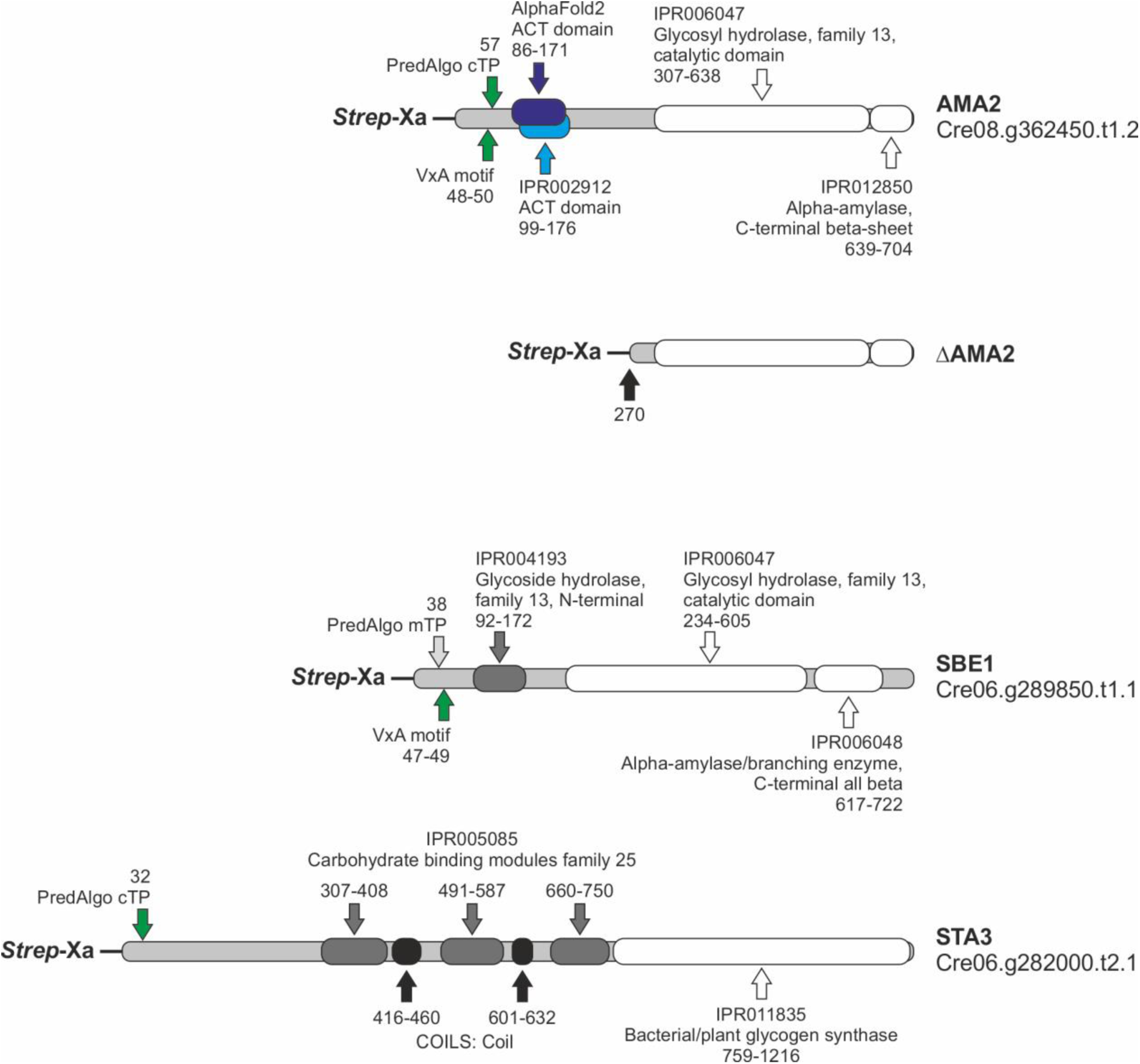
Schematic domain architectures of the analyzed enzymes. Domain architectures of the enzymes were analyzed by InterPro (Paysan-Lafosse et al., 2023). The AMA2 truncated variant ΔAMA2, described in the materials and methods section, is shown additionally. The primary sequences were retrieved from Phytozome v13, *Chlamydomonas reinhardtii* v5.6. InterPro entries (IPR) and descriptors as well as the positions of predicted domains are shown drawn to scale. PredAlgo (Tardif et al., 2012) transit peptide (TP) predictions as well as the presence of VxA motifs, typical for algal chloroplast TPs (Franzén et al., 1990; Tardif et al., 2012) are indicated. AMA2 was additionally modeled by AlphaFold (Supporting Fig. S6), and the ACT domain as predicted by the model is indicated in addition to the InterPro predictions. **cTP** / **mTP**: chloroplast / mitochondrial transit peptide.

**Supporting Figure S2.**
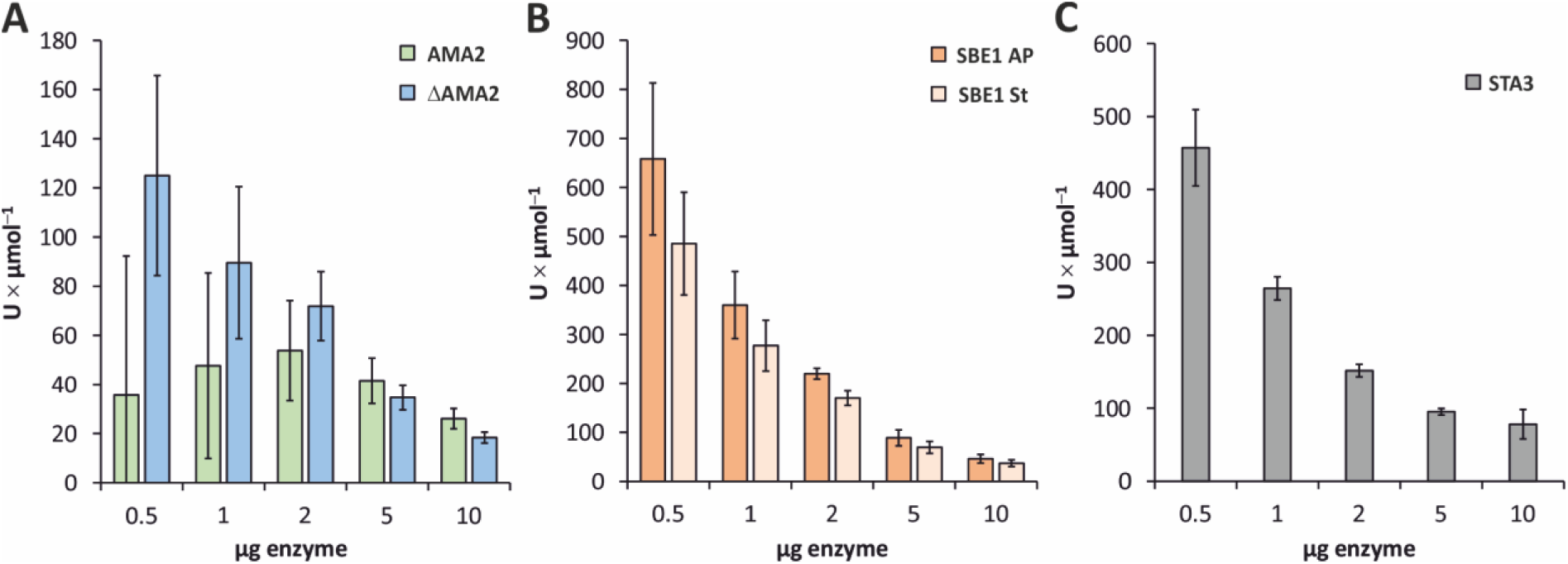
Determination of optimal enzyme amounts. *In vitro* activity assays were conducted with different amounts of AMA2, its amylase-domain-only variant ΔAMA2 (both in **A**), SBE1 (**B**) and STA3 (**C**) to identify optimal enzyme concentrations. Amylase and starch branching enzyme activities were analyzed by recording the absorbance of iodine-stainable soluble potato starch (**St** in **B**) and, in case of SBE1, amylopectin (**AP** in **B**). **A** and **B**: The indicated enzyme amounts, present in in 200 µL of 0.1 M potassium phosphate buffer, pH 7, or ‘physiological buffer’ (see the materials and methods section) were mixed with soluble potato starch or amylopectin and incubated at 30°C for 30 min. After quenching the reaction by adding HCl, the solutions were mixed with iodine solution and the absorption was determined at λ = 580 nm (starch) or λ = 550 nm (amylopectin). Enzyme activity (U) was defined as the decrease of 1 mg of iodine-stainable substrate per minute. **C**: STA3 activity was determined according to Kulichikhin et al. (2016). The indicated amounts of STA3 were added to the first reaction mixture. U of soluble starch synthase was defined as the release of 1 µmol of ADP per minute. **A** to **C**: The columns show average values, error bars the standard deviation. All assays were conducted with at least two independent enzyme preparations in at least three independent experiments. In case of AMA2, four enzyme preparations were analyzed in at least five independent assays, but seven to test the activities of 0.5, 1 or 2 µg enzyme.

**Supporting Figure S3.**
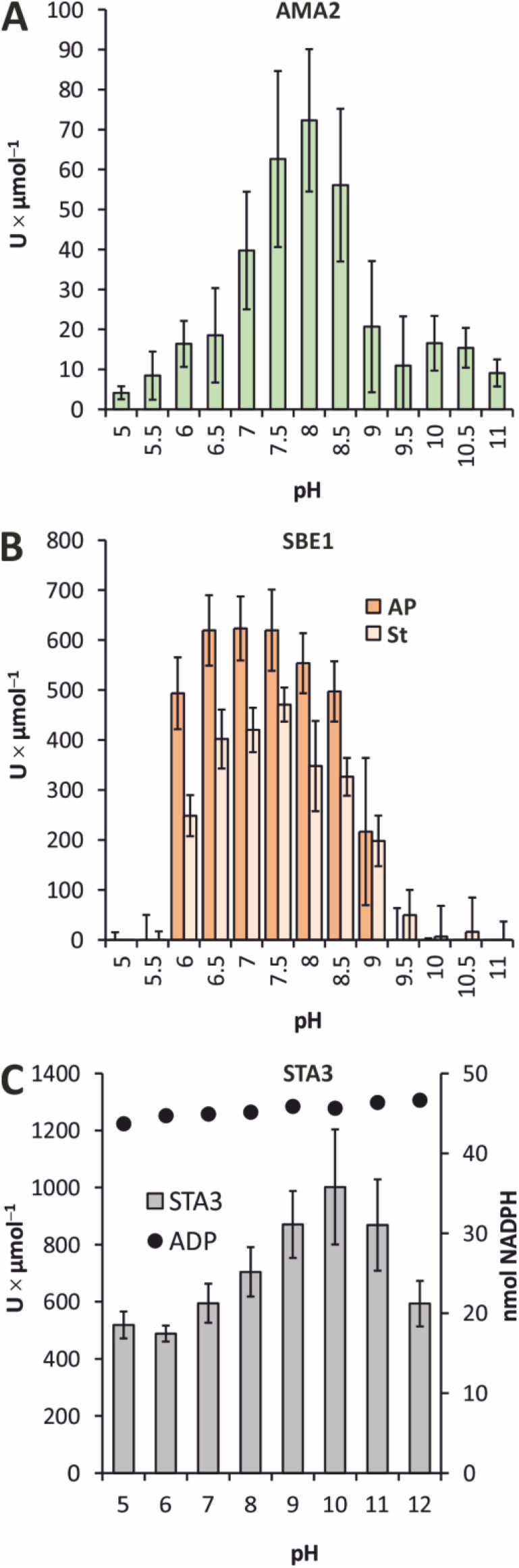
Determination of pH optima. **A** to **C**: Enzyme activity assays were conducted as described for Supporting Fig. S2, except that Britton-Robinson buffer, adjusted to the indicated pH values, was employed. Soluble potato starch (**St**) (**A**, **B**) or amylopectin (**AP**) (**B**) were used as substrates. **C**: In the case of STA3 (gray columns), the buffer of the first reaction mixture as described by Kulichikhin et al. (2016) was replaced by Britton-Robinson buffer of the indicated pH values. To test for a pH effect on the subsequent enzymes of this assay, controls were performed in which 100 µM ADP was added to first reaction mixture instead of STA3 for each pH value. The corresponding amounts of NADPH are indicated (black filled circles; secondary y-axis). **A** to **C**: All columns show average values, while standard deviations are indicated by the error bars. Three to four independent experiments were conducted in each case, employing two (SBE1), three (AMA2) or four (STA3) independent enzyme preparations.

**Supporting Figure S4.**
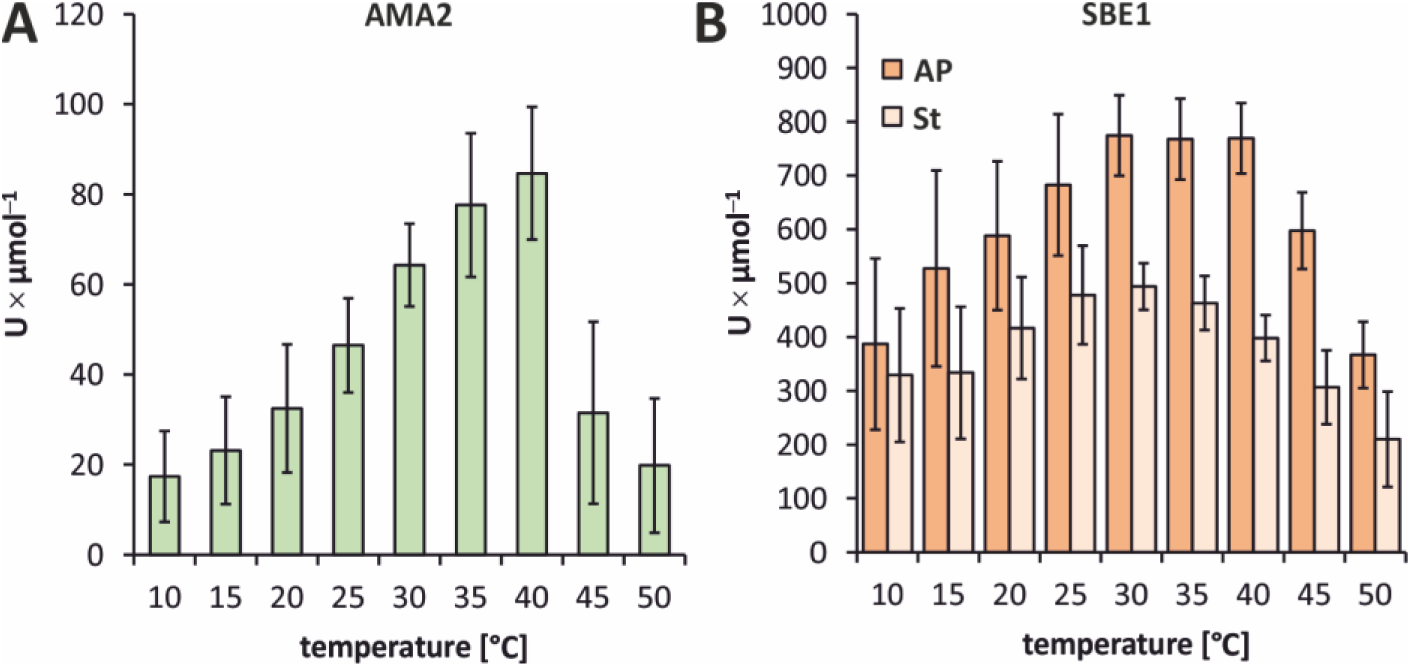
Determination of temperature optima. **A**, **B**: *In vitro* activity assays were conducted employing 2 µg (25.3 pmol) AMA2 (**A**) or 0.5 µg (5.7 pmol) of SBE1 (**B**). Amylase and starch branching enzyme activities were analyzed as described for Supporting Fig. S3 except that Britton-Robinson buffer pH 7.5 and pH 7 were used for AMA2 and SBE1, respectively, and that the incubation temperatures varied as indicated at the x-axis. SBE1 was assayed with both amylopectin (**AP**) and starch (**St**) as substrates. The assays were conducted with two independent enzyme preparations in four independent experiments. The columns show the average values, and the error bars show the standard deviations.

**Supporting Figure S5.**
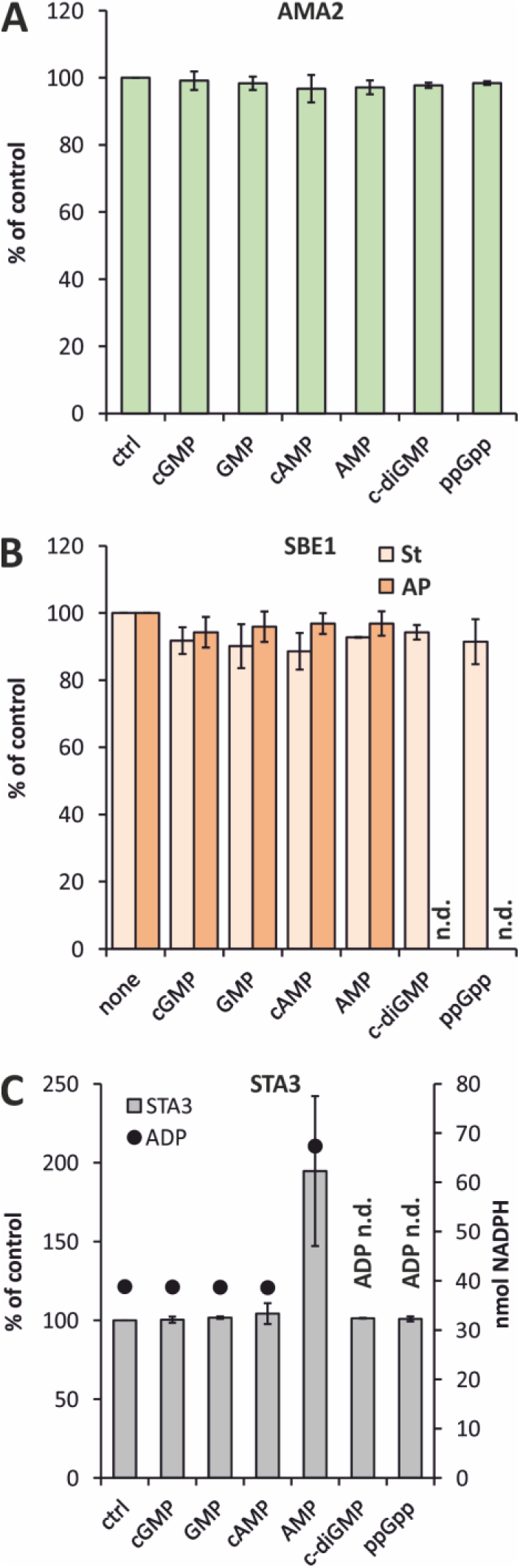
Testing the effects of nucleotides. **A** to **C**: The specific activities of 2 µg (25.3 pmol) AMA2 (**A**), 0.5 µg (5.7 pmol) SBE1 (**B**) or 0.5 µg (3.6 pmol) STA3 (**C**) were determined as described for Supporting Fig. S2, employing ‘physiological buffer’, pH 7.4, for AMA2 (**A**) and SBE1 (**B**). Cyclic guanosine or adenosine monophosphate (**cGMP**, **cAMP**), guanosine or adenosine monophosphate (**GMP**, **AMP**), cyclic di-GMP (**c-diGMP**) or guanosine tetraphosphate (**ppGpp**) were added directly to the reaction mixtures (only the first reaction mixture in case of STA3) to a final concentration of 100 µM. SBE1 activity was determined on both substrates amylopectin (**AP**) and starch (**St**) (**B**). The average values of the specific activities measured in technical triplicates (duplicates in the case of STA3) of the control (**ctrl**) reactions was set to 100 % for each independent experiment, and the specific activities determined in the presence of each nucleotide were set in relation to these. All assays were conducted with at least two independent enzyme preparations in at least two independent experiments. The columns show average values, and the error bars show the standard deviations. In case of the coupled-enzyme STA3 assay, the effect of nucleotides on the assay was tested once by replacing STA3 by 100 µM ADP (**C**; secondary y-axis). The specific activities determined in the control reactions were (**A**) 77.9 ± 11.8 U × µmol^-1 (^AMA2), (**B**) 579.5 ± 121.7 U × µmol^-1 (^SBE1, St) and 740.2 ± 30.9 U × µmol^-1 (^SBE1, AP), (**C**) 538.4 ± 90.1 U × µmol^-1 (^STA3).

**Supporting Figure S6.**
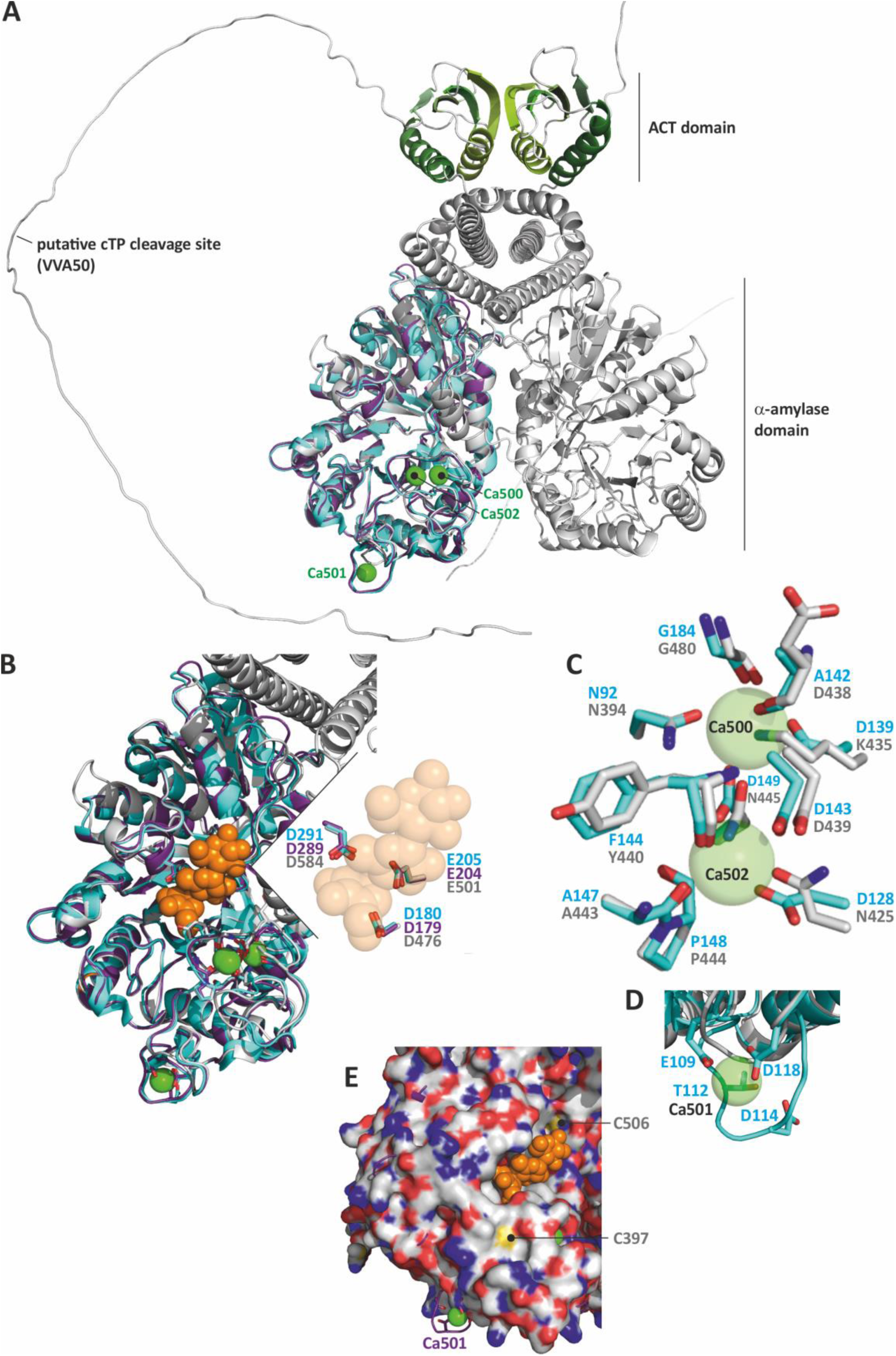
AlphaFold model of an AMA2 dimer and comparisons to Barley α-amylases 1 and -2. A model of the putative AMA2 dimer was generated by AlphaFold2-multimer and aligned with the crystal structures of Barley (*Hordeum vulgare*) α-amylases 1 and -2 (HvAMY1, HvAMY2) (HvAMY1: PDB: 1HT6, turquoise cartoons or carbon atoms; Robert et al. (2003)); HvAMY2 in complex with acarbose: PDB: 1BG9, purple cartoons or carbon atoms; Kadziola et al. (1998)). **A**: The structure model of AMA2 is shown as gray cartoon, except for the ACT domains, in which consecutive α-helices and β-strands were colored in shades of green. The AMA2 dimer model and the HvAMY2 structure (purple cartoon) were aligned to HvAMY1 (turquoise cartoon) in PyMOL. The Ca^2+ i^ons in the structures of HvAMY1 and HvAMY2 are shown as green spheres and labeled according to their numbers in the crystal structures (Robert et al., 2003, and references therein). The putative chloroplast targeting peptide (cTP) cleavage site of AMA2 is indicated at one of the termini (VAA50; 50 is the position within the AMA2 protein sequence Cre08.g362450.t1.2 including the N-terminal Met). **B**: Section of **A** that depicts the α-amylase domains only. The inhibitor acarbose within the structure of HvAMY2 is shown as orange spheres. The inset zooms into the active site and shows an overlay of the residues of the catalytic Asp-Glu-Asp triad. **C** and **D** show close-ups of the Ca^2+ i^ons and the coordinating residues of HvAMY1. The amino acids of AMA2 that correspond to those in HvAMY1 are overlain (also see the sequence alignment in Supporting Fig. S7). **E**: Overlay of AMA2 and HvAMY2, in which the surface is depicted for AMA2. Two Cys residues of AMA2 that are visible from the surface are indicated. **B** to **E**: Turquoise, purple and gray letters indicate residues and their numbers in the primary sequence of HvAMY1, HvAMY2 and AMA2, respectively, as aligned in Supporting Fig. S7.

**Supporting Figure S7.**
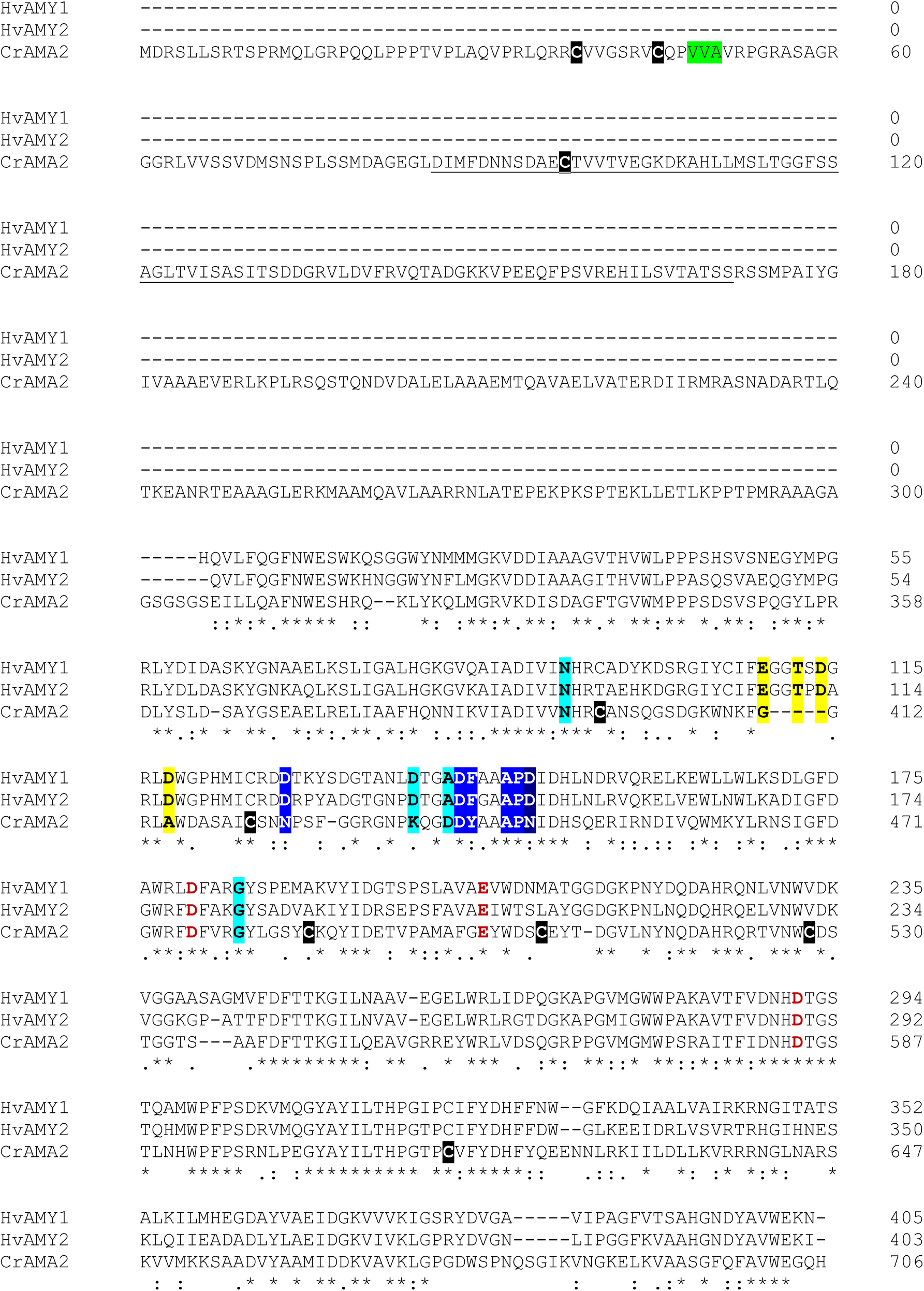
Alignment of AMA2 and Barley α-amylase sequences. The AMA2 primary sequence as indicated by gene model Cre08.g362450.t1.2 was aligned to the sequences linked to the crystal structures of Barley α-amylase 1 (abbreviated as HvAMY1; protein data bank (PDB) accession 1HT6) (Robert et al., 2003) and α-amylase 2 in complex with acarbose (HvAMY2; PDB accession 1BG9) (Kadziola et al., 1998) employing Clustal Omega. Asterisks below the alignment indicate fully conserved residues, while colons and periods show residues with strongly and weakly similar properties, respectively. For AMA2, the putative chloroplast targeting peptide cleavage site (VAA_50)_ is highlighted green, and Cys residues are written in white, bold letters that are highlighted black. The AMA2 sequence that forms the ACT domain as indicated by the AlphaFold model is underlined. The sequence alignment of HvAMY1 and HvAMY2 provided in Robert et al. (2003) as well as the structural overlays shown in Supporting Fig. S6 were employed as a guide to annotate the residues of the active site triad (bold red letters) and the residues forming Ca^2+-^binding sites in HvAMY1 and HvAMY2. Those that bind to Ca500, Ca501 and Ca502 (Ca^2+ n^umbering is according to Robert et al. (2003), and references therein) are shown here as bold letters highlighted turquoise, bold letters highlighted yellow, and white, bold letters highlighted blue, respectively. White, bold letters highlighted dark blue coordinate both Ca500 and Ca502.

**Supporting Figure S8.**
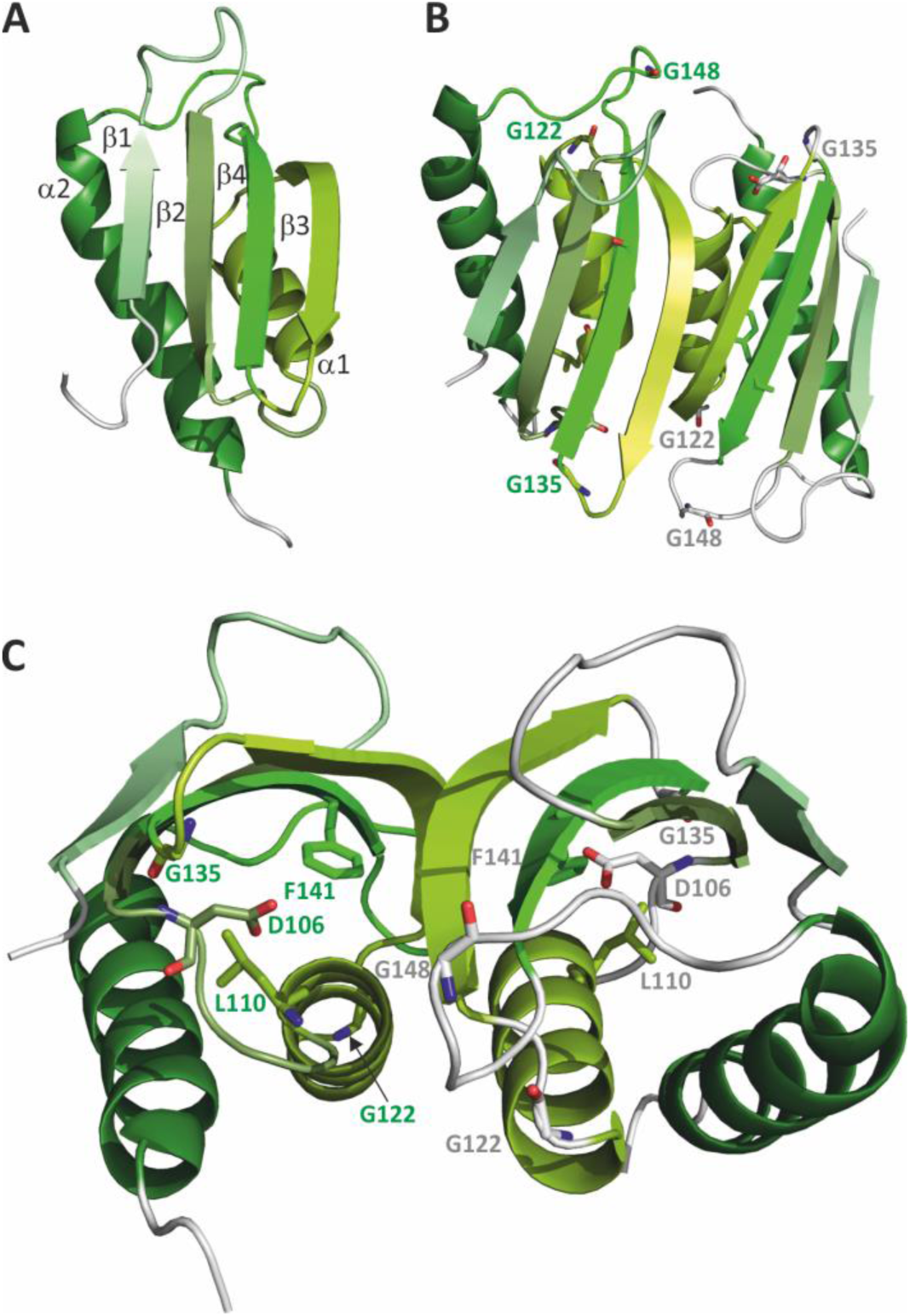
Model of the AMA2 ACT domain with exchanged residues labeled. The AMA2 ACT domain(s) depicted here are those of the AMA2 dimer model shown in Supporting Fig. S6A from which the other protein parts were hidden. **A**: Only one of the two AMA2 ACT domains is shown, and the α-helices and β-strands are numbered according to their order from N-to C-terminus. The β-strands are each labeled to their left. **B**: The dimeric arrangement of the AMA2 ACT domains is shown and the Gly residues that were exchanged in this study are labeled. **C**: The ACT domain dimer is shown from the side and the exchanged residues D106, L110 and F141 are indicated. The exchanged Gly residues shown in **B** are labeled when visible. **B** and **C**: Note that the loops of one monomer were colored in the same shades of green as their preceding α-helix or β-strand and the amino acid labels are written in green letters, whereas the loops were colored gray in the other monomer, and residues are labeled by gray letters.

**Supporting Figure S9.**
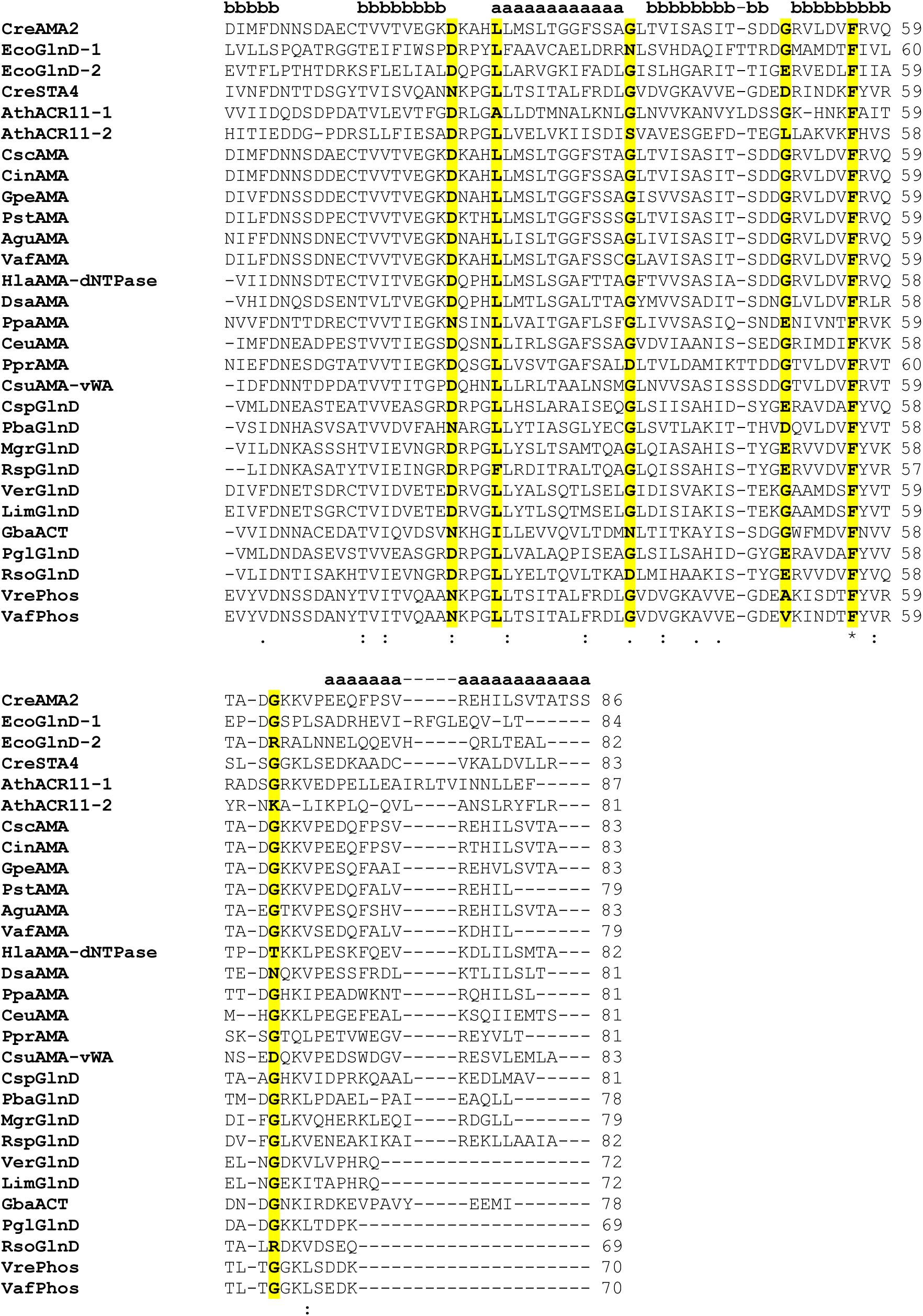
Alignment of ACT domain sequences. Sequences similar to the AMA2 ACT domain were retrieved employing NCBI’s BlastP tool and aligned using Clustal Omega after the manual addition of the ACT domain sequences of *E. coli* GlnD (**EcoGlnD**; UniProt B6HZE1), *Arabidopsis* ACR11 (**AthACR11**; The Arabidopsis Information Resource (TAIR) AT1G16880) and *Chlamydomonas* STA4 (**CreSTA4**; Cre12.g552200.t1.2). EcoGlnD and AthACR11 contain two ACT domains each, indicated by the suffixes -1 and -2. See the materials and methods section for more details on the proceeding and Supporting Table S1 for additional information on the aligned sequences. The presence of β-sheets (**b**) or α-helices (**a**) of the *Chlamydomonas* AMA2 ACT domain as predicted by AlphaFold are indicated on top. Residues that align to the amino acids exchanged in AMA2 are written in bold and highlighted yellow, independent from their degree of conservation, which is indicated by asterisks (fully conserved), colons and periods (amino acids with strongly and weakly similar properties, respectively). Abbreviations used are a combination of species name and protein domains predicted in addition to an ACT domain: **Species name abbreviations: Cre**: *Chlamydomonas reinhardtii*, **Eco**: *Escherichia coli*, **Ath**: *Arabidopsis thaliana*, **Csc**: *Chlamydomonas schloesseri*, **Cin**: *Chlamydomonas incerta*, **Gpe**: *Gonium pectorale*, **Pst**: *Pleodorina starrii*, **Agu**: *Astrephomene gubernaculifera*, **Vaf**: *Volvox africanus*, **Hla**: *Haematococcus lacustris*, **Dsa**: *Dunaliella salina*, **Ppa**: *Polytomella parva*, **Ceu**: *Chlamydomonas eustigma*, **Ppr**: *Pycnococcus provasolii*, **Csu**: *Coccomyxa subellipsoidea* C-169, **Csp**: *Caulobacter* sp., **Pba**: Planctomycetota bacterium, **Mgr**: *Magnetospirillum gryphiswaldense*, **Rsp**: Rhodospirillales bacterium, **Ver**: Verrucomicrobia bacterium, **Lim**: Limisphaerales bacterium, **Gba**: *Gossypium barbadense*, **Pgl**: *Phenylobacterium glaciei*, **Rso**: *Rhodovibrio sodomensis*, **Vre**: *Volvox reticuliferus*. **Domain abbreviations: AMA**: alpha amylase, **dNTPase**: p-loop containing nucleoside triphosphate hydrolase, **vWF**: von Willebrand factor type A domain, **GlnD**: [protein-PII] uridylyltransferase/uridylyl-removing enzyme, **ACT**: ACT domain-only (plant ACR proteins), **Phos**: starch / glycogen phosphorylase.

**Supporting Figure S10.**
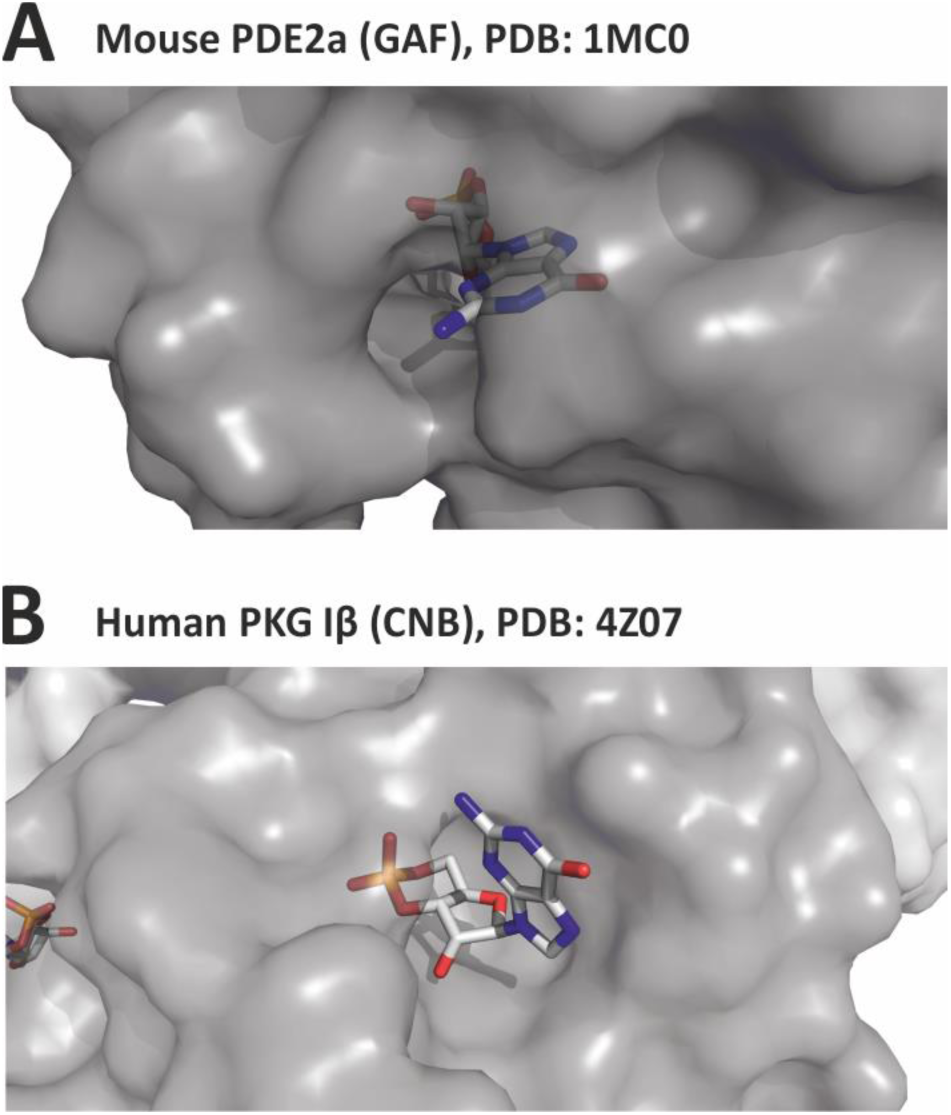
Exemplary cGMP-binding modes. Examples for cGMP bound to a GAF-(cGMP-dependent phosphodiesterases (PDEs), adenylyl cyclases, and FhlA)-(**A**) and a CNB-(Cyclic Nucleotide-Binding)-(**B**) domain. Both figures were prepared in PyMOL, showing the protein surface in transparent gray. The cGMP molecules are shown as sticks, and colored by elements (gray: C, blue: N, red: O, orange: P). **A**: GAF domain of mouse PDE2a, PDB: 1MC0 (Martinez et al., 2002). **B**: CNB domain of human protein kinase Iβ, PDB: 4Z07 (Kim et al., 2016).

## Notes

### Competing Interest Statement

The authors have declared no competing interest.

